# Systematic assessment of blood-borne microRNAs highlights molecular profiles of endurance sport and carbohydrate uptake

**DOI:** 10.1101/721928

**Authors:** Fabian Kern, Nicole Ludwig, Christina Backes, Esther Maldener, Tobias Fehlmann, Artur Suleymanov, Eckart Meese, Anne Hecksteden, Andreas Keller, Tim Meyer

## Abstract

Multiple studies endorsed the positive effect of regular exercising on mental and physical health. However, the molecular mechanisms underlying training-induced fitness in combination with personal life-style remain largely unexplored. Circulating biomarkers such as microRNAs (miRNAs) offer themselves for studying systemic and cellular changes since they can be collected from the bloodstream in a low-invasive manner. In *Homo sapiens* miRNAs are known to regulate a substantial number of protein-coding genes in a post-transcriptional manner and hence are of great interest to understand differential gene expression profiles, offering a cost-effective mechanism to study molecular training adaption, and connecting the dots from genomics to observed phenotypes.

Here, we investigated molecular expression patterns of 2, 549 miRNAs in whole-blood samples from 23 healthy and untrained adult participants of a cross-over study, consisting of 8 weeks of endurance training, with several sessions per week, followed by 8 weeks of washout and another 8 weeks of running, using microarrays. Participants were randomly assigned to one of the two study groups, one of which administered carbohydrates before each session in the first training period, switching the treatment group for the second training period. During running sessions clinical parameters as heartbeat frequency were recorded. This information was extended with four measurements of maximum oxygen uptake (VO2 max) for each participant.

We observed that multiple circulating miRNAs show expression changes after endurance training, leveraging the capability to separate the blood samples by training status. To this end, we demonstrate that most of the variance in miRNA expression can be explained by both common and known biological and technical factors. Our findings highlight six distinct clusters of miRNAs, each exhibiting an oscillating expression profile across the four study timepoints, that can effectively be utilized to predict phenotypic VO_2_ max levels. In addition, we identified miR-532-5p as a candidate marker to determine personal alterations in physical training performance on a case-by-case analysis taking the influence of a carbohydrate-rich nutrition into account. In literature, miR-532-5p is known as a common down-regulated miRNA in diabetes and obesity, possibly providing a molecular link between cellular homeostasis, personal fitness levels, and health in aging.

We conclude that circulating miRNAs expression can be altered due to regular endurance training, independent of the carbohydrate availability in the timeframe around training. Further validation studies are required to confirm the role of exercise-affected miRNAs and the extraordinary function of miR-532-5p in modulating the metabolic response to a high availability of glucose.

## 1 Introduction

The positive effects of sports activity on physical and mental health as well as the cardiovascular effects of training have been widely characterized [1, 2, 3]. In contrast, the current understanding on how the genetic information is related to sports activity and how molecular processes are affected by exercise is still limited. Earlier studies showed that physical exercise has an impact on epigenetic factors, which are closely related to aging, such as DNA methylation levels, or the composition of histone modifications [4, 5]. For example, Ludlow *et al* demonstrated that exercising activates certain genes responsible for repairing DNA damage, in addition to the MAPK signalling pathway that ultimately contributes to telomere stability [6]. Also, expression levels of non-coding RNA transcripts in particular microRNAs (miRNAs) were found to be associated with exercising [7, 8]. MiRNAs, typically 22 nucleotides in length, are loaded into proteins of the AGO family and orchestrate post-transcriptional gene regulation by targeting mainly 3^*1*^ UTRs of partially complementary mRNAs [9]. Circulating miRNAs, i.e. molecules carried by the bloodstream, constitute low-invasive biomarkers that can be extracted from whole-blood, serum, or plasma samples [10]. Previous work successfully utilized blood-borne miRNA profils to identify adaptive molecular mechanisms as triggered by physical activities [11, 12]. This motivates the question whether miRNA signatures being indicative of fitness levels after repeated endurance exercise, commonly measured via increasing values of maximal oxygen consumption VO_2_ max, exist [12].

Here, we analyzed 90 blood samples using microarrays to determine miRNA expression patterns from a randomized cross-over study, in particular to investigate the molecular relation between endurance sports and changes in VO_2_ max [13]. Moreover, the influence of variations in carbohydrate-based nutrition on VO_2_ max levels was examined by searching for miRNAs that can differentiate between participants that show a positive training effect in combination with glucose uptake before each session and those that should refrain from any administration.

## 2 Results

### 2.1 Outline

Our findings are based on a randomized cross-over study that consists of 23 participants, randomly split into two groups. All participants were both healthy and untrained as indicated by a medical check including history taking, physical examination, resting and exercise ECG. The cohorts are *N*_1_ = 13, *N*_2_ = 10 consisting of 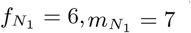, and 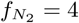 females, 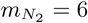 males, respectively. During study conduction participants were between 30 and 62 years old. Throughout the course of the study each participant completed 8 continuous weeks with 4 × 45 minutes endurance training sessions per week, followed by a wash-out phase, and finally another 8 weeks of training analogous to the first interval. Directly before and after each training interval, a blood-sample was taken from each participant, resulting in four timepoint measurements *E*_1_, *A*_1_, *E*_2_, and *A*_2_. In addition, during the first training interval participants that were assigned to the second group consumed 50 g glucose monohydrate dissolved in water, 15 minutes before each training session. Likewise, participants from the first group applied a carbohydrate solution before sessions of the second training phase (Figure 1). Furthermore, several anthropometric and fitness parameters such as the ventilatory threshold (VT1), maximum oxygen uptake (VO_2_ max), weight and Body mass index (BMI), body fat levels, and maximal heart rate frequency were recorded (Supplementary Table 1) as well as a complete blood count (CBC) for each sample. After preparation full-blood samples were hybridized using a microarray to measure the expression of 2,549 human miRNAs from miRBase release v21 [14]. Overall, 90 of 92 samples could be measured successfully. Further, 307 miRNAs remained after removing those exhibiting either a low detection rate or a low-expression distribution across the samples. More details are provided in the methods section. While for the average sample Spearman correlation is at or above 80%, an outlier sample could be identified that was subsequently excluded, leaving 89 samples for in-depth analysis (Supplementary Figure 1, top row and leftmost column). Also, no evident technical and biological batch effects could be found that predominantly influenced the miRNA-sample matrix clustering (Supplementary Figure 2). However, there is a trend in both matrix clustering approaches for the samples to be grouped by the *Timepoint* of blood extraction, notably not by each single timepoint individually but in a pairwise manner of pre-training (*E*_1_, *E*_2_) and post-training (*A*_1_, *A*_2_) timepoints, as indicated by the dichotomous variable *Training state*.

**Figure 1:**
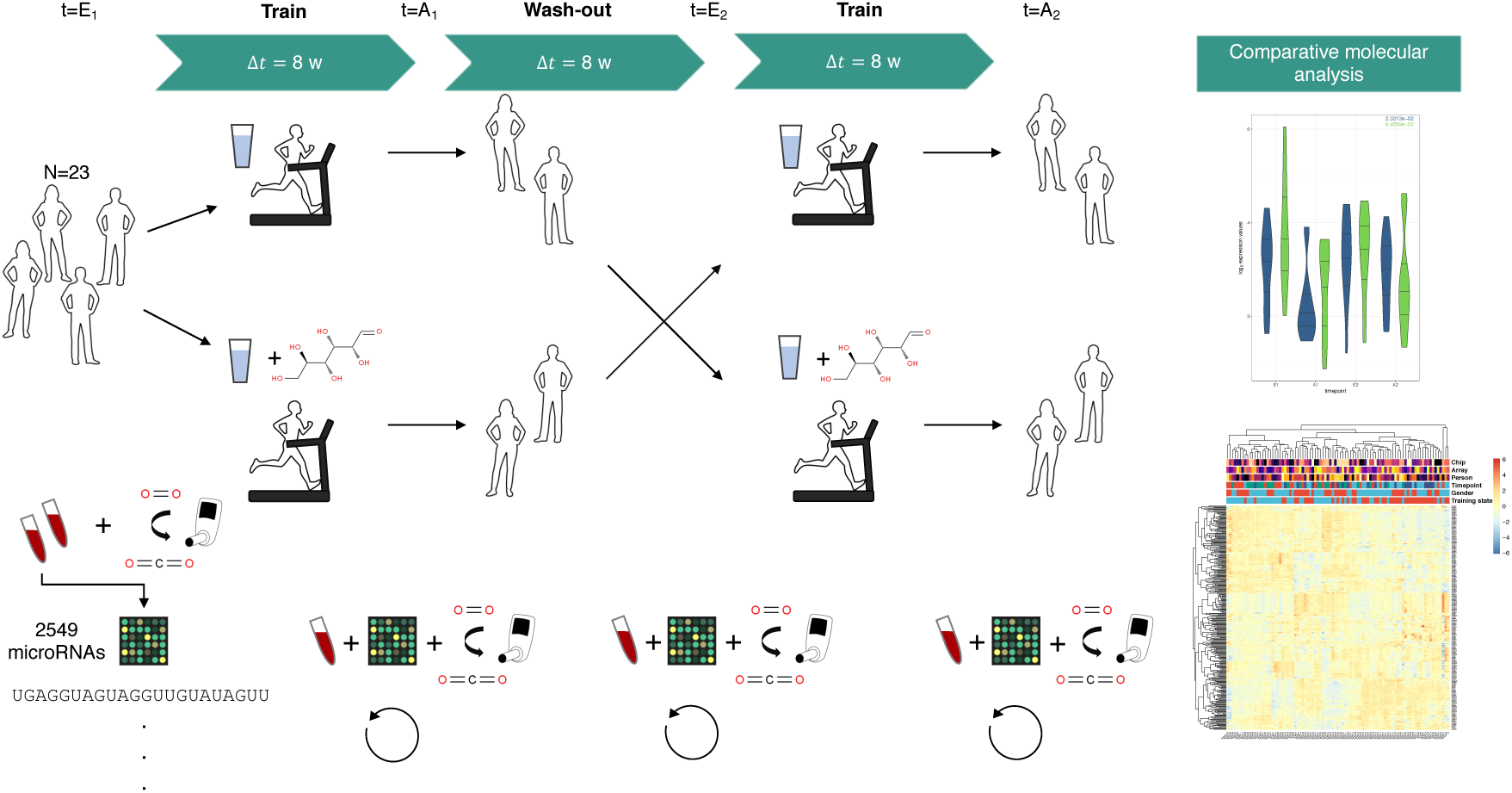
Overview on study design. Healthy and untrained participants were randomly assigned to any of two training groups, each performing of 8 weeks of 4 × 45 min training, followed by a wash-out phase, again followed by 8 weeks of endurance training. In the first period participants of one group orally administered glucose-solution 15 minutes before each running session, while participants of the second group administered carbohydrates in their second training period (cross-over). At four timepoints (*E*_1_, *A*_1_, *E*_2_, and *A*_2_) a blood-sample was taken and measured using cDNA-Microarrays probed with 2, 549 human microRNAs.

**Figure 2:**
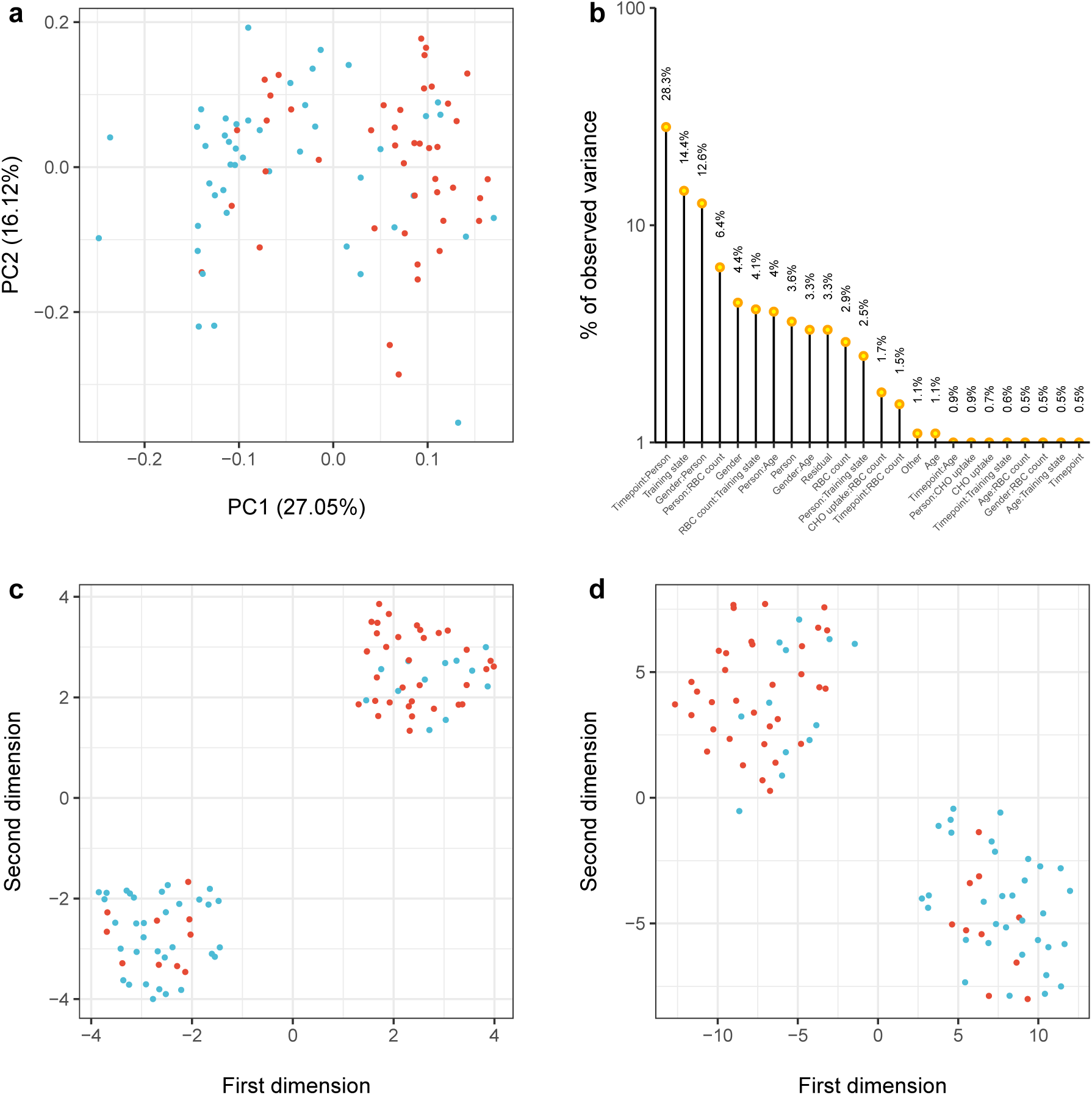
Analysis of variance and factors explaining it using measured miRNA expression data. **a** Sample distribution within the first two principal components obtained from PCA along with the percentage of variance explained in each dimension. **b** Results from Principal Variance Component Analysis (PVCA) showing estimates of variance in the expression data that can be explained with both known and unknown (hidden) sample annotation factors. Each bar corresponds to one factor, where mixed interactions between two variables are also possible and marked respectively by a colon. **c** Two-dimensional UMAP embedding using the miRNA-sample matrix *X* ∈ ℝ^90×307^. **d** Two-dimensional t-SNE embedding using the miRNA-sample matrix *X*. In each dimension reduction panel a single point corresponds to one sample that is colored according to the timepoint of blood extraction and relative to the training period, i.e. before (blue) or after (red) one of the training intervals.

### 2.2 Physical exercising affects microRNA expression

To investigate whether endurance training is reflected by a molecular change of miRNA expression we analyzed the samples using both methods for dimension reduction and classical differential expression analysis. A two-dimensional embedding of the samples using Principal Components Analysis (PCA) and the results for a corresponding batch variate assessment with Principal Variance Component Analysis (PVCA) are shown in panel **a** and **b** of Figure 2 [15, 16]. In total, the first two principal components account for approximately 43% variance in the miRNA-sample matrix. As visible in panel (**a**) samples measured before a training interval are enriched in the halfspace for which *PC*1 ≤ 0.02. Conversely, post-training samples are enriched in the halfspace *PC*2 > 0.02. Inspection of the sub-spaces spanned by the eigenvectors PC1 and PC2 did not yield an extreme distribution of prominent key-features but revealed a set of 65 miRNAs with a loading coefficient larger than 0.1 (Supplementary Table 2). Interestingly, the largest factor explaining the observed variance in miRNA expression is the interaction between timepoint and person connoted with each sample, suggesting that miRNAs show a substantial inter-individual expression in healthy individuals. Further, the second most informative variable corresponds to whether a participant was untrained, i.e. the sample was taken before any training interval, or trained, i.e. the sample was taken after the 8-week training intervals. Apparently, the gender of an individual seems to play a role as well, as it is estimated to explain nearly 13% variance. Since it is known that endurance training influences the abundance of red blood cells (RBCs), which make up the largest portion of cells within full-blood samples, we assessed whether a potential increase in RBC counts confounds the power of the variable *Training state* in separating the samples. Even though a certain part of the observed variance in miRNA expression can be justified with a changing number of RBCs, it is certainly not the most prominent factor. Remarkably, information whether the sample was taken after a training interval in which a participant administered carbohydrates as indicated by the surrogate variable *CHO uptake* substantiates only a minor fraction in our analysis. To overcome the inherent linear dependencies as uncovered through PCA, the more complex dimension reduction methods Uniform Manifold Approximation and Projection (UMAP) and t-Distributed Stochastic Neighbor Embedding (t-SNE) both further improved the separation of pre- and post-training samples, suggesting that non-linear effects play a role in our data set as well [17, 18]. Because a similar number of the sample annotations disagree with the observed over-representation of *Training state* in the two clusters, we investigated whether the samples belong to a specific set of participants that show a differential expression pattern, which turned out not to be the case. Indeed, both clusters show a significant enrichment according to a hypergeometric test with respect to the colored *Training state* variable (*P* ≈ 4.3 × 10^−6^ for Cluster 1 located top-left, *P* ≈ 1.4 × 10^−6^ for Cluster 2 in bottom-right corner of panel **d**, Figure 2).

Next, we assessed to which extent miRNAs are differentially expressed taking a pooled sample approach, once neglecting the CHO treatment groups, and once taking the different courses of training into account. Respective volcano plots for the different comparison setups are shown in Figure 3. After pooling samples from both groups (setup 1) for the first training interval (*E*_1_, *A*_2_), 11 miRNAs are significantly down-regulated after 8 weeks of endurance training with a fold-change ≤ −1 (miR-378a-5p, miR-223-3p, miR-30b-5p, miR-199a-5p, miR-7977, miR-326, miR-30c-5p, miR-550a-3p, miR-331-3p, miR-484, and miR-340-3p). Moreover, 13 miRNAs significantly increased in expression (let-7g-5p, miR-144-5p, miR-98-5p, let-7a-5p, miR-17-3p, miR-126-3p, miR-195-5p, miR-20b-5p, miR-20a-5p, let-7f-5p, miR-26b-5p, miR-199a-3p, and miR-144-3p). Consulting the miRNA PCA-loadings for the two sets confirms their informative importance, each showing a larger mean of loadings than compared to the overall set of features (*mean*_*down*_ ≈ 0.099, *mean*_*up*_ ≈ 0.099, and *mean*_*all*_ ≈ 0.076). MiRNA enrichment analysis on the set of down-regulated genes has been done using an online tool that computes which biochemical functional categories are significantly enriched in miRNA gene sets or lists (MiEAA). The MIEAA analysis revealed among others the protein-coding target gene APLN (*p* ≈ 0.048, *FDR* < 0.05) known to be associated with cardiovascular homeostasis and glucose metabolism [19, 20]. For the group specific comparison, we found that 22 miRNAs are up-regulated (let-7g-5p, miR-18b-5p, miR-454-3p, miR-374a-5p, miR-18a-5p, miR-16-5p, miR-93-5p, miR-98-5p, let-7a-5p, miR-7107-5p, miR-17-3p, miR-126-3p, miR-15a-5p, miR-195-5p, miR-20b-5p, miR-374b-5p, miR-20a-5p, miR-17-5p, let-7f-5p, miR-26b-5p, miR-199a-3p, and miR-144-3p) and 10 are down-regulated (miR-30a-5p, miR-378a-5p, miR-30b-5p, miR-326, miR-30c-5p, miR-550a-3p, miR-331-3p, miR-16-2-3p, miR-484, and miR-30d-5p) in samples of participants that orally administered a glucose solution before each training session (Figure 3, panel(**c**)). In contrast, considering expression levels from participants that did not ingest a glucose solution 9 miRNAs are up-regulated (let-7g-5p, miR-144-5p, miR-98-5p, miR-17-3p, miR-1246, let-7f-5p, miR-26b-5p, miR-199a-3p, and miR-144-3p) while 10 show decreased expression (miR-378a-5p, miR-223-3p, miR-30b-5p, miR-199a-5p, miR-7977, miR-192-5p, miR-30c-5p, miR-550a-3p, miR-484, miR-340-3p) (Figure 3, panel(**e**)). Even though no single mature miRNA exhibits opposed expression patterns between the two groups, the observed changes in differentially expressed miRNAs remain significant (*p* ≈ 4.8 × 10^−24^, fisher’s exact test). Surprisingly, miR-144-3p is the only feature that is down-regulated using pooled samples of the second training interval (Figure 3, panel(**b**)), denoting that no significant alterations for the CHO-treatment and group specific comparisons can be reported.

**Figure 3:**
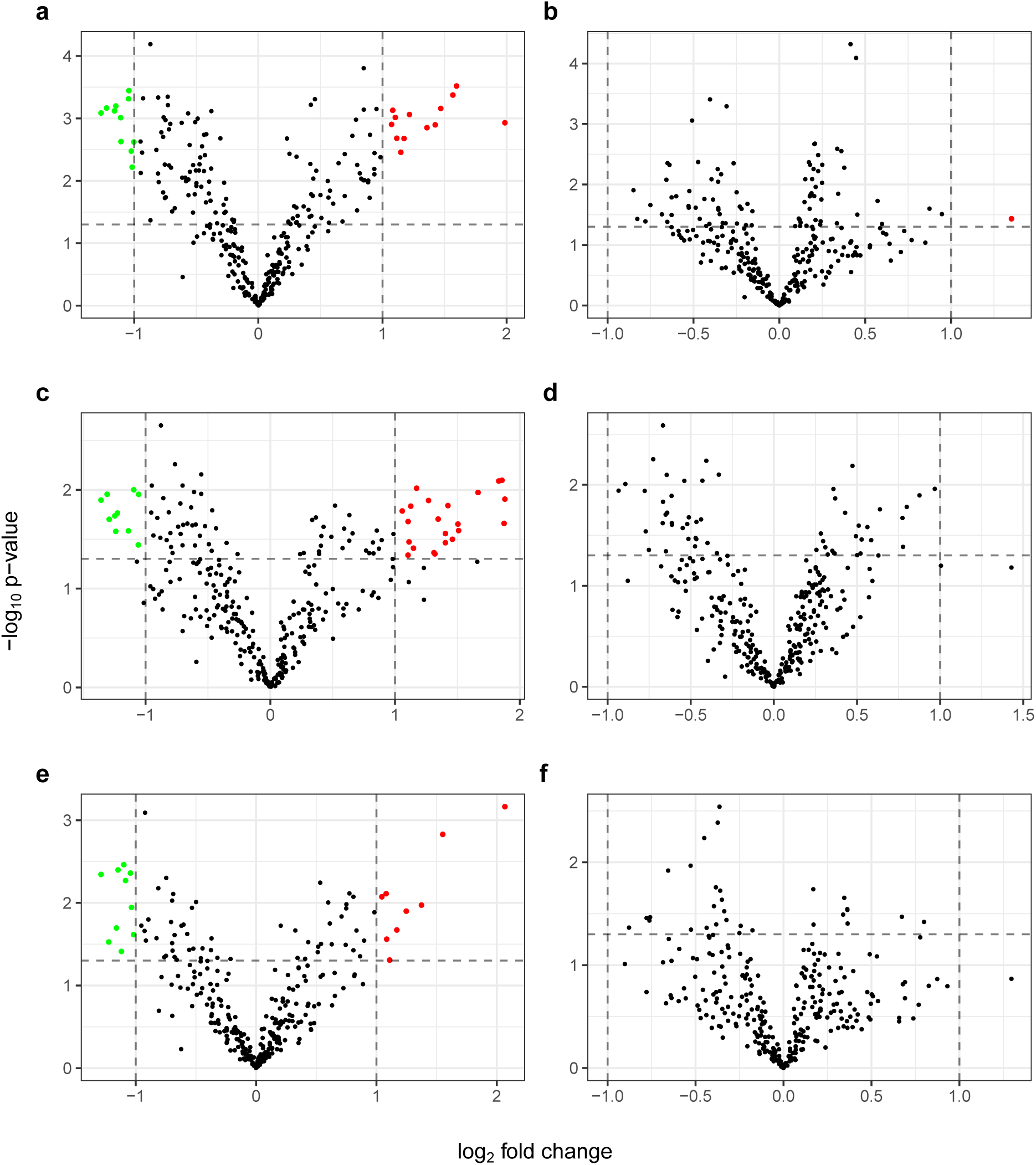
Volcano plots for six comparison setups using 307 microRNAs. The x-axis indicates log_2_ fold change, while the y-axis marks the negative decade logarithm of p-values from student’s t-tests (unadjusted). Dashed horizontal lines indicate a p-value of 0.05, while dashed vertical lines indicate a fold change of 2. **a** Pooling the samples from the two participant groups and using timepoints of the first training interval (*E*_1_ and *A*_1_). **b** Pooled sample approach analogous to **a** but with timepoints from the second training period (*E*_2_ and *A*_2_). **c** Analysis for timepoints like in **a** only using samples from the first treatment group. **d** Analysis for timepoints like in **b** only using samples from the first treatment group. **e** Analysis for timepoints like in **a** only using samples from the second treatment group. **f** Analysis for timepoints like in **b** only using samples from the second treatment group. MiRNAs that exceed both axis thresholds (dashed lines) are colored according to their direction of dys-regulation, i.e. red for up-regulation and green for down-regulation after training.

Taking into account our findings about the differential expression of miRNAs we investigated if certain groups of miRNAs exist that exhibit similar expression patterns and whose expression is consistently changed after training. Using the full z-score scaled miRNA-sample expression matrix six miRNA expression clusters 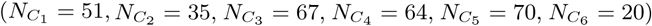 were highlighted (Figure 4). The computed disjoint miRNA to cluster assignments are listed in Supplementary Table 3. Broadly speaking, the clusters can be separated into two larger expression classes, each showing a wave-like expression pattern with either a positive z-score peak first (*C*_1_, *C*_3_, and *C*_5_) or a negative z-score peak first (*C*_2_, *C*_4_, and *C*_6_). Notably, the direction of expression change is mostly maintained within in each cluster across the two training intervals, where the first peak, which corresponds to the first round of exercising, in general appears to be more prominent than the second, also reflected by the volcano plots in Figure 3. Nevertheless, differences between the clusters can also be observed, since *C*_1_, *C*_3_, and *C*_6_ are much more attenuated than the remaining ones. These findings not only provide an explanation for the good separability of the samples with respect to trained and untrained participants, but propose that physical exercising may have diverse consequences on the molecular level and that a subset of the human miRNome is consistently affected by physical exercising.

**Figure 4:**
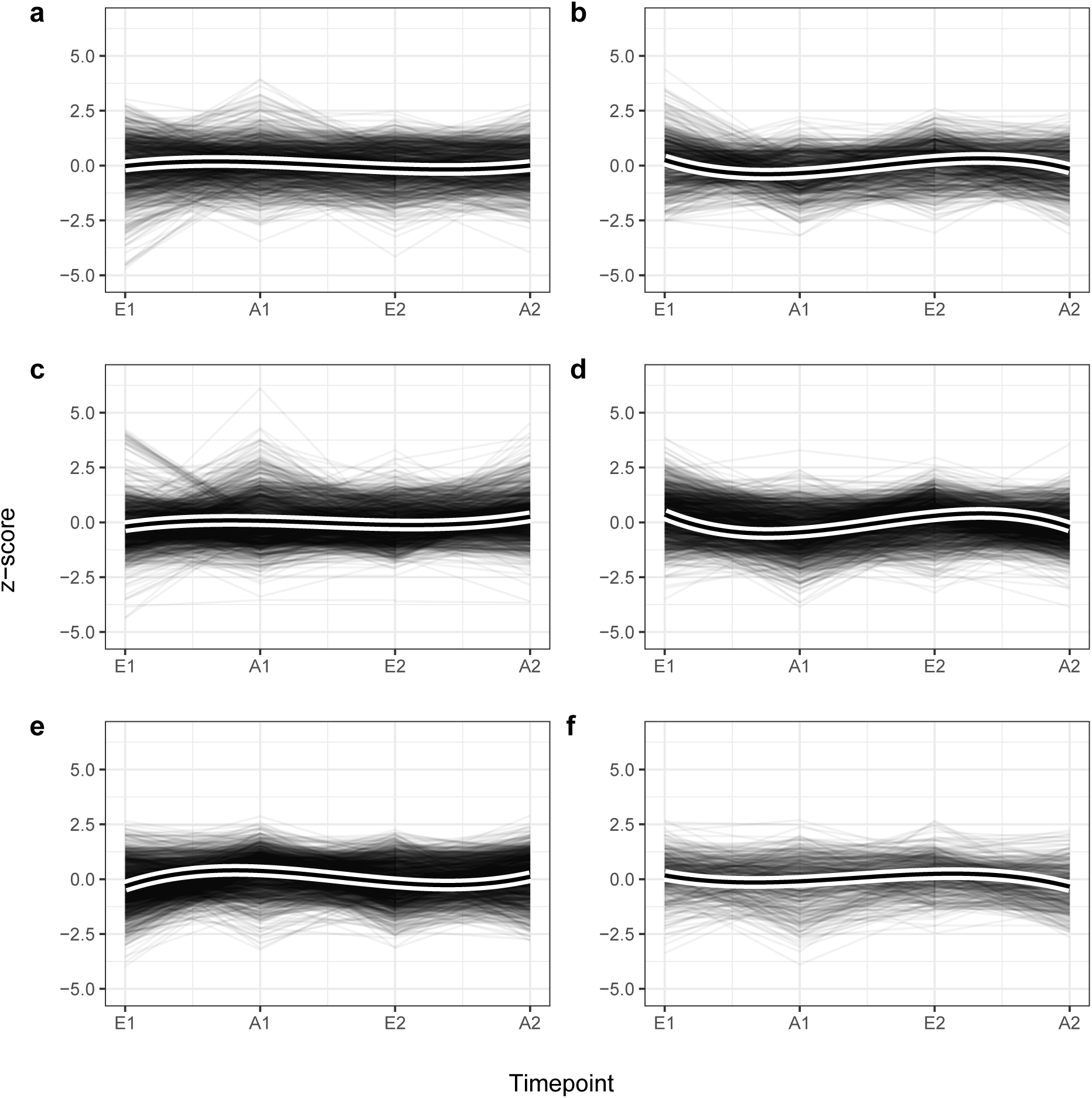
Distribution of Z-scores divided by six miRNA clusters *C*_1_ - *C*_6_ corresponding to panel **a** - **f**, along the four study timepoints using all samples. Each single grey line corresponds to one miRNA expression profile for one participant. Thick black lines display cluster-specific and smoothed curves of a cubic b-spline basis (*b* = 3). The distinct clusters contain microRNAs that exhibit similar expression patterns over time. Although every cluster exhibits a wave-like expression pattern, miRNAs in *C*_1_, *C*_3_, *C*_5_ show a tendency to be up-regulated after training (*up, down, up*) while those in *C*_2_, *C*_4_, and *C*_6_ tend to be down-regulated after a period of endurance exercise (*down, up, down*).

### 2.3 MiRNA expression levels correlate with change in VO_2_ max

Because our results indicate that miRNAs can be affected by endurance training, we put the analysis a step further and hypothesized that expression levels correlate with phenotypic changes in training performance observed through VO_2_ max. It is widely accepted that an improvement in VO_2_ max levels after endurance training denotes an improvement in the personal functional capacity. To this end, we computed the Spearman correlation between measurements VO_2_ max and miRNA expression values for each participant (Supplementary Figure 3). In the resulting correlation matrix three clusters of participants can be recognized, each showing an enrichment towards the group the participant were assigned to, i.e. CHO first and Non-CHO second versus Non-CHO first and CHO second training interval. Interestingly, several miRNAs show an opposing correlation towards VO_2_ max in-between the participant clusters, whereas some seem not to show any robust correlation pattern.

To pinpoint potential candidate miRNAs that exhibit a good correlation with the observed phenotype we performed linear regression using the miRNA expression values as features and the VO_2_ max values as dependent variable. Overall the model reached an estimated *R*^2^ ≈ 0.41, suggesting that miRNAs partially explain the variance in VO_2_ max, yet questioning the existence of other factors that remain to be explored. In Figure 5 the distribution of the top 20 positive and top 20 negative feature coefficients from the best model selected with repeated cross-validation is shown. A link back to the expression groups reveals a dissimilar distribution of cluster identities between miRNAs with a different sign of coefficient. While positive coefficients are distributed among all clusters, cluster 1, 2, and 6 are mostly depleted in the top list of negative feature coefficients. This result affirms a putative correlation between exercise-induced, oscillating miRNA expression profiles and altered levels in VO_2_ max. In total 86 out of 307 miRNAs received a linear coefficient unequal from zero. Also, we report the absolute feature importance of each variable with a non-zero coefficient using the trained model (Supplementary Table 4). To check whether the miRNAs selected by our model are known to be associated with important molecular functions, we used MiEAA to perform over-representation analysis. Among the pathways, organs, and biological functions enriched for the list of miRNAs having a non-zero feature importance we identified besides others; *Upregulated in male* (*P* ≈ 3.7 × 10^−4^, *FDR* < 0.05), *AMPK signaling* (*P* ≈ 2.6 × 10^−3^, *FDR* < 0.05, Wikipathways: WP1403), and *Insulin signaling pathway* (*P* ≈ 1.4 × 10^−4^, *FDR* < 0.05, KEGG: hsa04910).

**Figure 5:**
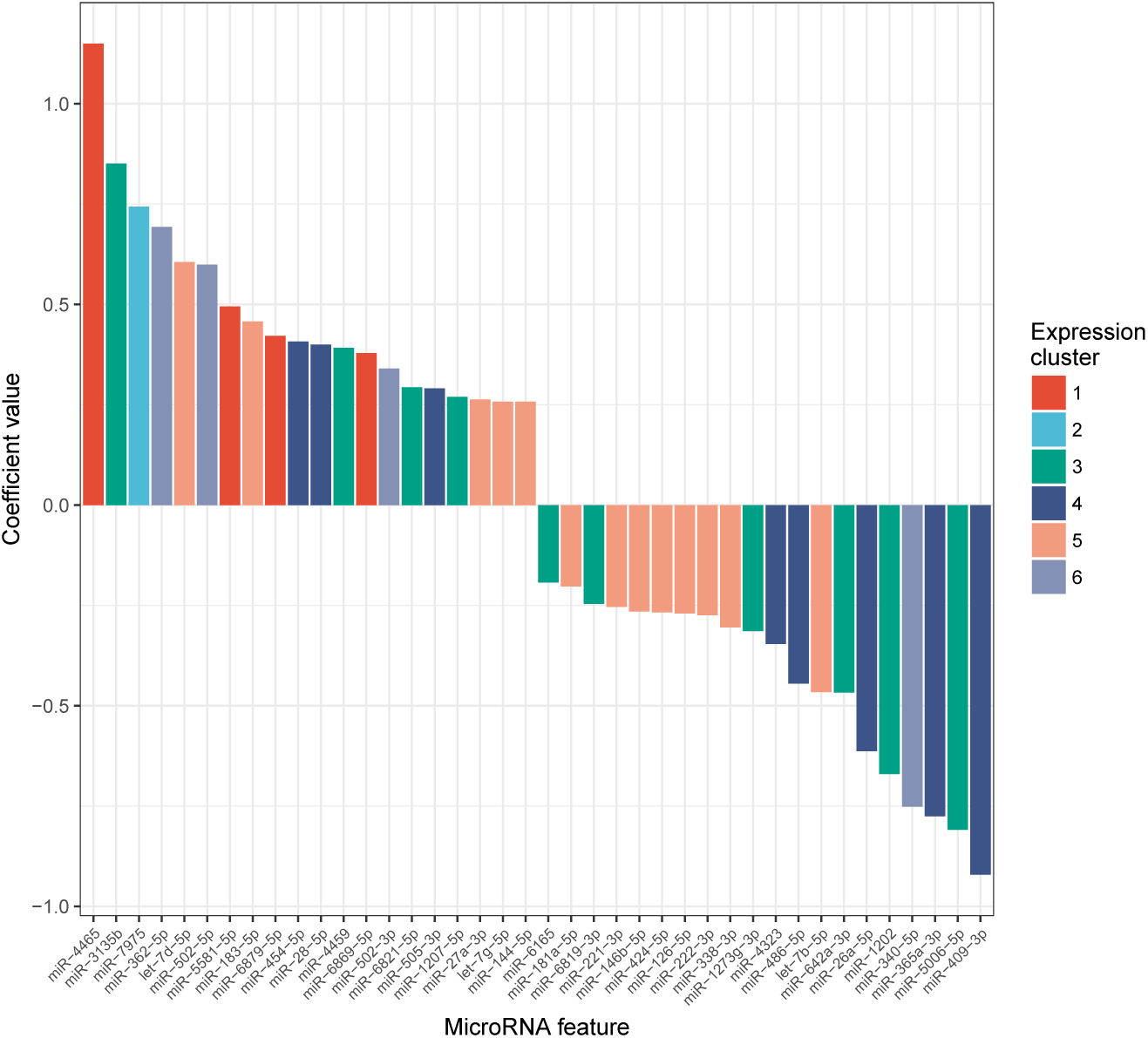
Top 20 positive and negative miRNA regression coefficients. Feature coefficients stem from the best linear model with respect to *R*^2^ predicting the measured VO_2_ max values. In total 86 out of 307 miRNAs were assigned a coefficient unequal from zero. Each miRNA bar is colored according to its cluster identity (*C*_1_ - *C*_6_) from Figure 4, highlighting a differential distribution of these clusters among the most important regression coefficients.

### 2.4 Candidate marker miR-532-5p indicates change in VO_2_ max after carbohydrate uptake

Previous studies about the effect of a carbohydrate-rich nutrition on the training performance, in particular an impairment of the homeostasis as conferred by AMPK, yielded conflicting results. Therefore, we state the alternative hypothesis that this effect is highly individual, depending on many intra-personal factors such as sleep habits, age, gender, and genetic factors. Hence, we asked whether the observed correlations between miRNA and VO_2_ max levels can be leveraged to separate the cohort in two recommendation groups, one that either showed a greater change in VO_2_ max in positive direction, i.e. greater improvement, or a smaller change in negative direction, i.e. less worsening under CHO treatment and the second group defined as the complement of group 1, namely participants for which the conditions apply under non-treatment periods. The grouping process is summarized and displayed as decision tree in panel **a** of Figure 6. Using the above outlined decision process we could separate the participants into two almost balanced groups (CHO recommended: *N*_1_ = 10, CHO not recommended: *N*_1_ = 9). Consequently, we assessed the miRNA expression levels for each timepoint, additionally taking into account the two recommendation groups. Manual inspection of 307 distributions highlighted a differential expression pattern of miR-532-5p, shown in panel **b** of Figure 6. In general, miR-532-5p is clearly expressed above background and potentially is affected by endurance training since both mean expression and variance increase for samples measured after any of the two training periods, independent of the recommendation assignments. First, the difference in pre-training samples for which no carbohydrates were administered was not significant between the two groups (*P* ≈ 0.15, Welch Two Sample t-test for *T*_1_ dark red *vs. T*_1_ dark blue, panel **b** of Figure 6, left), while the post-training difference turns out to be significant (*P* ≈ 0.024, Welch Two Sample t-test for *T*_2_ light red *vs. T*_2_ light blue, panel **b** of Figure 6, left). Surprisingly, when repeating the analysis for the same recommendation groups but taking only the CHO treatment intervals into account, the differences vanish. Not only the pre-training difference remains to be insignificant (*P* ≈ 0.51, Welch Two Sample t-test for *T*_1_ dark red *vs. T*_1_ dark blue, panel **b** of Figure 6, right) but also the post-training difference gets insignificant (*P* ≈ 0.99, Welch Two Sample t-test for *T*_2_ light red *vs. T*_2_ light blue, panel **b** of Figure 6, right). Following candidate selection, we investigated how miR-532-5p is expressed in-between the recommendation groups closely in relation to observed levels of VO_2_ max. To this end, we propose to assess the potential of miR-532-5p as a molecular marker in order to judge a possible effect of CHO uptake and a subsequent response in training performance by only considering the non-treatment period for each participant. We observed not only that the Spearman correlation between log_2_ fold change of miR-532-5p and change in VO_2_ max turns sign (*ρ* ≈ −0.4 pos. recommendation, *ρ* ≈ 0.26 neg. recommendation) but miR-532-5p being even more up-regulated in the positive recommendation group (Figure 6, panel **b**, left). Also, according to our definitions the mean change in VO_2_ max is lower as compared to the negative recommendation group. Conversely, for samples from the individual treatment periods, the Spearman correlation for the first group switches accordingly, while for the second it approximately remains the same. Hence, our results suggest that exercised-induced up-regulation of miR-532-5p is associated with observed levels of VO_2_ max and that it has potential discriminatory power in separating individuals who might administer carbohydrates before training from those that should refrain from a carbohydrate-rich nutrition.

**Figure 6:**
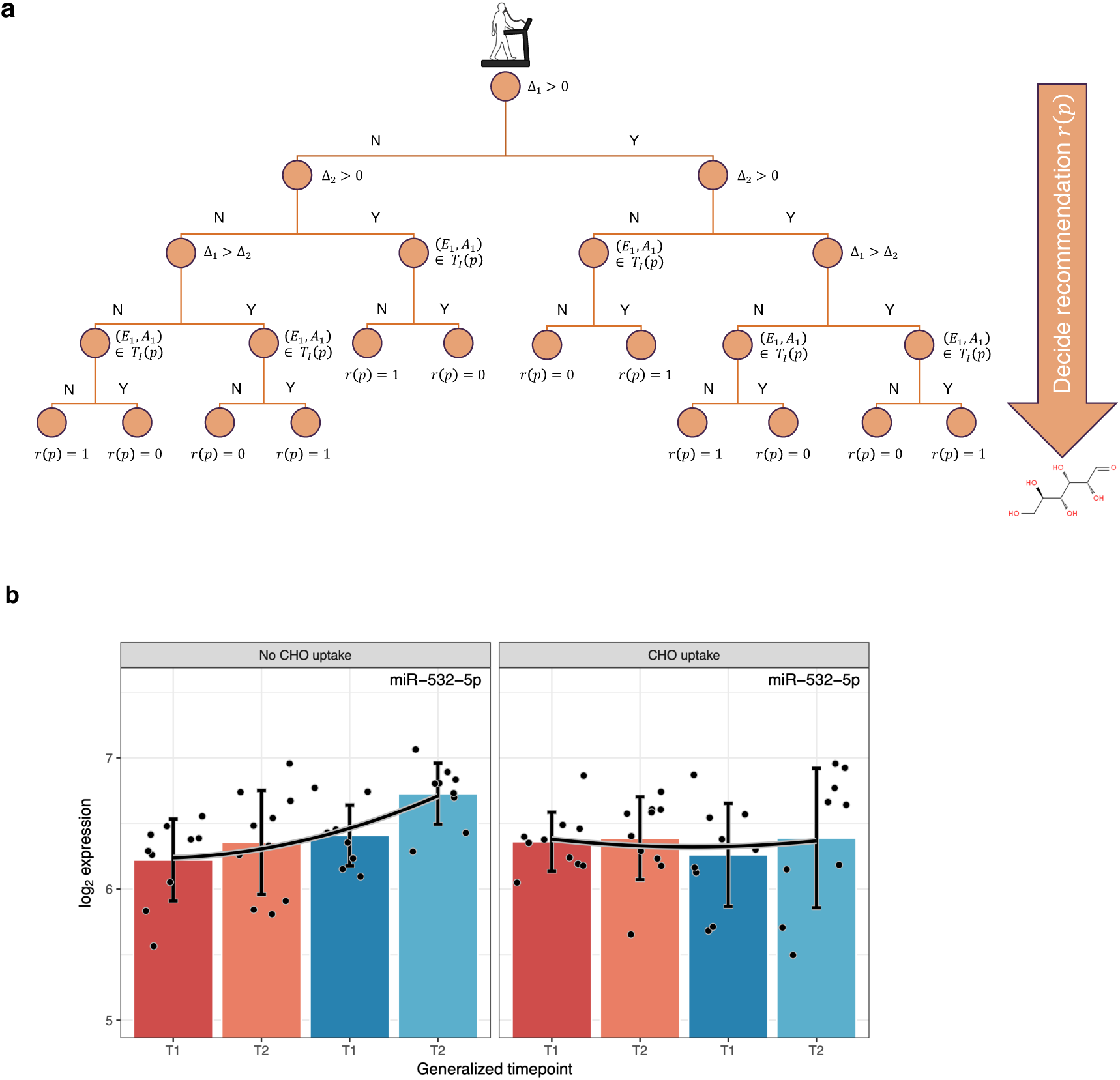
Assignment procedure conducted to find CHO recommendation groups and group-wise expression of candidate marker miR-532-5p. **a** Decision tree for devised procedure to classify each participant *p* into one of the two recommendation groups *r*(*p*) ∈ {0, 1}. The decision process primarily depends on the personal change in VO_2_ max values during the first training period, Δ_1_ = *V O*_2_(*A*_1_) − *V O*_2_(*E*_1_) and Δ_2_ = *V O*_2_(*A*_2_) − *VO*_2_(*E*_2_) the second training period. Further, *T*_*I*_ (*p*) describes the set of pairs of timepoints, i.e. the rounds during which the carbohydrates were administered. **b** Paired barplots showing the expression of miR-532-5p across training period timepoints, i.e. *T*1 ∈ {*E*_1_, *E*_2_} and *T*2 ∈ {*E*_2_, *A*_2_}, compared between the two CHO treatment recommendation groups, once using training periods without and once with an oral administration of glucose. Recommendation groups are highlighted by distinct colors where blue corresponds to a positive and red to a negative recommendation on glucose uptake. Each black point belongs to one sample in the distribution. Bar heights display the mean expression values and black error bars mean *±* standard deviation. Smoothed quadratic b-splines (*b* = 2) are drawn as black curves with a contrast margin.

## 3 Discussion

The prospective role of miRNAs as circulating biomarkers in diagnostics, which can be sampled in a low-invasive manner, has been already described by numerous publications [21, 22, 23]. However, technical factors such as sampling protocols, library preparations, sequencing technology, and statistical data analysis need to be fixed in order to account for potential bias and to make studies comparable among each other [24, 25]. Here, we analyzed four timepoint measurements of 23 participants from a random cross-over study to investigate the effect of endurance-training on blood miRNA expression in combination with a variation of the glucose availability during training sessions. Even though one cannot rule out all possible influences of bias, especially in small-scale studies, the stringent study design and rigorous data analysis suggests that multiple miRNAs are differentially regulated after 8 weeks of regular endurance training. This is reflected by our dimension reduction analysis where two clearly distinguishable clusters of samples after 8 weeks of regular endurance training with differing CHO supply prior to each session can be seen that, to the best of our knowledge, cannot be fully explained by any observed variable but most appropriately by the binary training state. Interestingly, the number of miRNAs with significantly changed expression dropped substantially after the second training period, an observation for which we propose carry-over effects to be the main reason, also due to the fact that participants were completely untrained at the beginning of the study [13].

Further, we observed a correlation between molecular changes in miRNA expression and phenotypic changes in VO_2_ max. Even though our miRNA enrichment analysis showed significantly associated biological categories such as the *Insulin signaling pathway*, the set of important miRNAs was rather large, possibly because some features are correlated among each other, motivating the need for further validation studies to increase specificity of the candidate set. Also, we suggest that inter-individual correlation profiles make up a large portion of variance in the data making it difficult to generalize the observed interrelations. Regarding miR-532-5p as a candidate marker for training performance outcome after carbohydrate uptake, an independent study by Cui *et al*. found that miR-532-5p was de-regulated in participants conducting muscular strength endurance training, and it showed a negative correlation with the insulin-like growth factor-1 [26]. Moreover, several earlier publications investigated the prominent role of miR-532-5p in the context of adult obesity and diabetes type 2, in particular insulin resistance [27, 28, 29, 30]. Interestingly, most of these studies have in common that miR-532-5p is down-regulated in patients suffering from obesity or diabetes. Setting these reports into context, our findings imply that miR-532-5p is up-regulated in individuals that exhibit a gain in training performance while consuming glucose before each training. This highlights a potential role of miR-532-5p in understanding homeostasis-related cell fitness as well as metabolic and systemic diseases, e.g. diabetes. It is important to note that the miR-532 precursor is located on the gonosomal X chromosome. Therefore, our results might be affected by a certain degree of gender bias due to a genetic imbalance of genomic locis. However, the gender distributions of our recommendation groups were not particularly skewed towards either female or male participants.

Taken together, our results imply a connection between genetic and molecular factors, i.e. miRNAs, and trainability of healthy human individuals, which might be characterized in a precise manner with the aid of larger validation studies.

## 4 Materials and Methods

### 4.1 Study design

Our findings are based on a randomized cross-over study in 23 healthy, previously untrained adults [13]. In brief, two eight-week training periods were separated by a wash-out period of equally eight weeks. Training consisted of 4 times 45 min running / walking at 70% of heart rate reserve per week. Compliance was supervised and documented by supervised training sessions and read-out of heart rate monitors. Each participant received 50 g of glucose dissolved in 200 ml of water 15 min before every training session of one of the two training periods (N1: glucose during second training period; N2: glucose during first training period; N1 = 13, N2 = 10 consisting of fN1 = 6, mN1 = 7, and of fN2 = 4 females, mN2 = 6 males, respectively). VO2 max was determined by an exhaustive, ramp-shaped exercise test with gas exchange measurements before and after each period. Blood was collected before the exercise tests. The original study design, methods and protocols are in accordance with the Declaration of Helsinki and approved by the local ethics committee (Ärztekammer des Saarlandes, Saarbrücken, Germany; approval number: 148/10). Written informed consent was obtained from all participants before exercise testing. This trial was registered at clinicaltrials.gov as NCT02297646.

### 4.2 Blood extraction

Whole Blood was collected in PAXGene Blood RNA Tubes (BD Biosciences). Total RNA including small RNAs were isolated using PAXGene Blood miRNA kit (Qiagen) according to manufacturers recommendations. RNA quantity and quality was checked with Nanodrop2000 (Thermo Fisher Scientific) and Bioanalyzer RNA 6000 Nano Kit (Agilent), respectively.

### 4.3 Microarray experiments

MiRNA profile was measured with Agilent Human miRNA microarray (miRbase v21) according to the manufacturers protocol. In short, 100ng total RNA was dephosphorylated and 3’ labelled with 3-pCp. Labelled RNA was hybridized to the array for 20 hours at 55°C an 20 rpm. Slides were washed and air-dried and subsequently scanned with Agilent Microarray Scanner (G2505C) with 3*µ*m resolution in double-path mode. Signals were extracted using Agilent Feature extraction software (version10.10.1.1).

### 4.4 Statistical analysis

Raw microarray data resulting from the Agilent array-scanner was parsed into a raw expression table and microRNA expression levels for the samples under consideration were quantile normalized and log_2_ transformed. We required a microRNA to be detected in at least 50% of the samples in order to be considered for subsequent analysis. Then, we discarded all microRNAs for which the 3rd quantile of log_2_ transformed expression values was below 3.5. Using sample Spearman correlation analysis we identified one outlier (Sample no. 26) that was removed from the dataset before performing any tests. All data analysis was performed using the statistical programming language R v3.5.3. Expression and correlation heatmaps were created using the package pheatmap v1.10.12. All other main and supplementary Figures were generated using the packages ggplot2 v2.2.1, ggfortify v0.4.6, ggsci v2.9, cowplot v0.9.4, grid v3.5.3, gridExtra v2.3. Common data manipulations such as merging, filtering, and sorting of tables were accomplished with data.table v1.12.2, reshape2 v1.4.3, dplyr v0.8.1, stringr v1.4.0, tidyr v0.8.3, purrr v0.3.2, and tibble v2.1.1. Principal components analysis was taken out using the *prcomp* function, in addition to the Principal Variants Component Analysis that was taken from the package pvca v1.20.0, having fixed the percentage of variance explained threshold at 90%. The t-SNE analysis was done with Rtsne v0.15 using the following parameters: dims: 2, initial dims: 50, perplexity: 20, theta: 0.0, check duplicates: TRUE, pca: FALSE, normalize: FALSE and keeping the rest at default, while the UMAP analysis was performed with umap v0.2.2.0 and the parameters: method: “naive”, n neighbors: 20, n components: 2, metric: “euclidean”, n epochs: 500, and min dist: 0.1, also keeping all other parameters at default. P-values are adjusted with *FDR* < 0.05 unless stated otherwise in the main text. Output of R builtin base functions such as t.test was transformed to data.frames using the broom package v0.5.2. Machine learning procedures were performed with caret v6.0-84. For the microRNA-VO_2_ max regression model we used repeated cross-validation with 10 repeats and 6 folds each. As data model glmnet with a tunelength of 50 was applied to find the best hyperparameters. Variable feature importance is defined as the absolute coefficient value resulting from the trained linear regression model.

### 4.5 Data availability and Accession codes

Microarray data is available through NCBI’s Gene Expression Omnibus (GEO) using the Accession ID GSE133910.

## 5 Conclusions

Further and independent cross-over studies are required in order to prove the role of the miRNA profiles and the candidate marker miR-532-5p as described in this study. A more encompassing characterization of the human blood-borne miRNome through next-generation-sequencing (sncRNA-seq) might reveal novel hypotheses and genetic variants that could be related to an endurance-training driven change in VO_2_ max.

## Author contributions

conceptualization, F.K., A.H., A.K., and T.M.; methodology, F.K., C.B., T.F., and A.S.; software, F.K., C.B., and A.S.; validation, N.L., C.B., A.H., A.K., and T.M.; formal analysis, F.K., C.B., and A.K.; investigation, F.K., N.L., E.M., and A.H.; resources, N.L., A.H., and A.K.; data curation, F.K., N.L., C.B., and A.H.; writing–original draft preparation, F.K.; writing–review and editing, F.K., N.L., C.B., T.F., A.H., A.K., and T.M.; visualization, F.K. and A.K.; supervision, A.K. and T.M.; project administration, A.K.; funding acquisition, A.K.

## Acknowledgements

The authors thank all participants of the original study for their voluntary contributions and efforts to stick to the stringent training conditions.

## Abbrevations

The following abbreviations are used in this manuscript:

VO_2_: max Maximal aerobic capacity
miRNA: microRNA
CHO: Carbohydrates

