## Supplementary document for "Systematic assessment of blood-borne microRNAs highlights molecular profiles of endurance sport and carbohydrate uptake"

Supplementary data

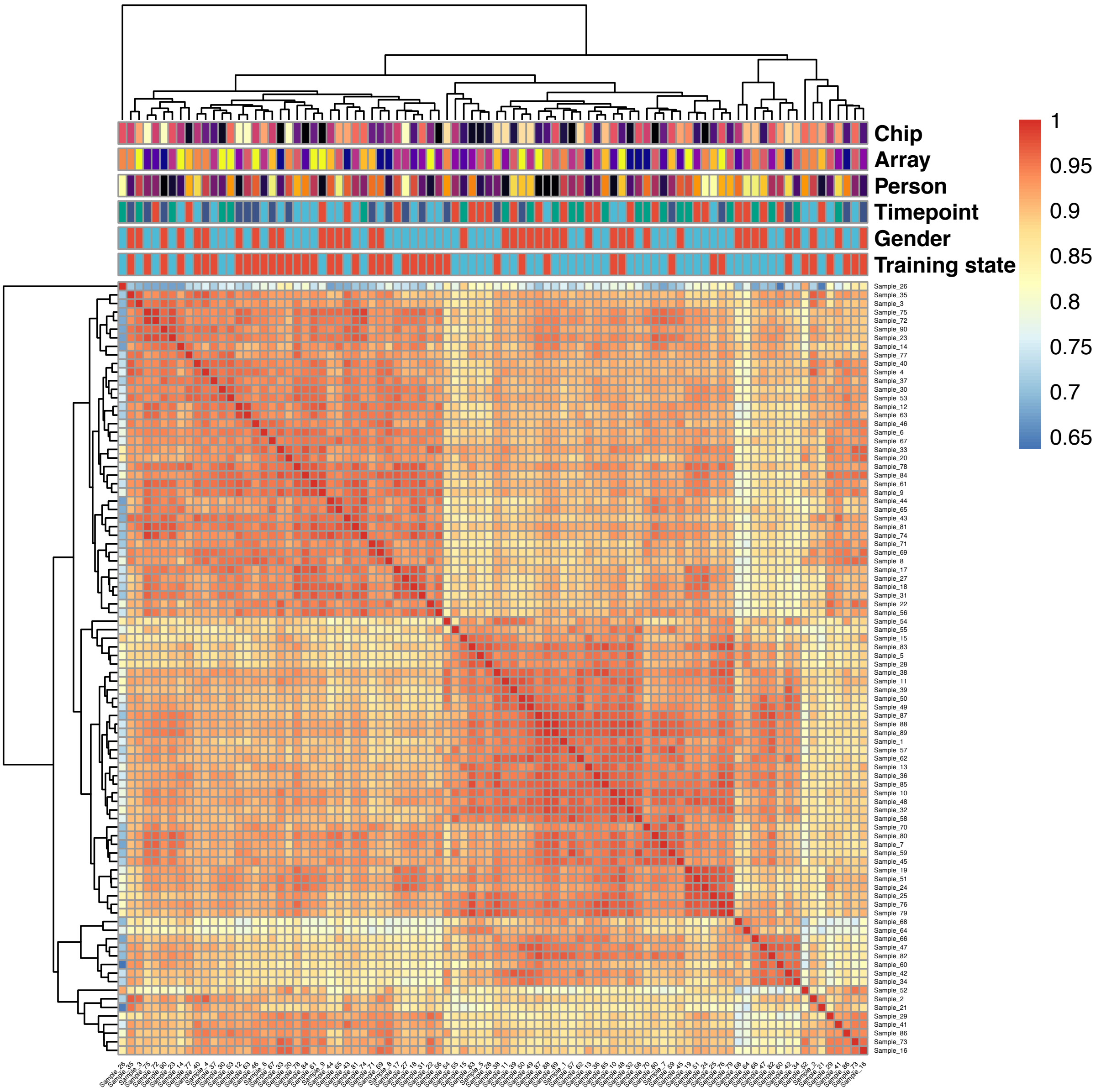


**Figure S1:** Sample to sample Spearman correlation matrix. Color coding according to correlation values where a higher correlation is shown in red, while lower values are shown in blue. Along each dimension a hierarchical clustering is plotted as a dendrogram. Color bars on top of the matrix encode for different annotation variables for each of the samples. The top and leftmost sample no. 200 was identified as an outlier.

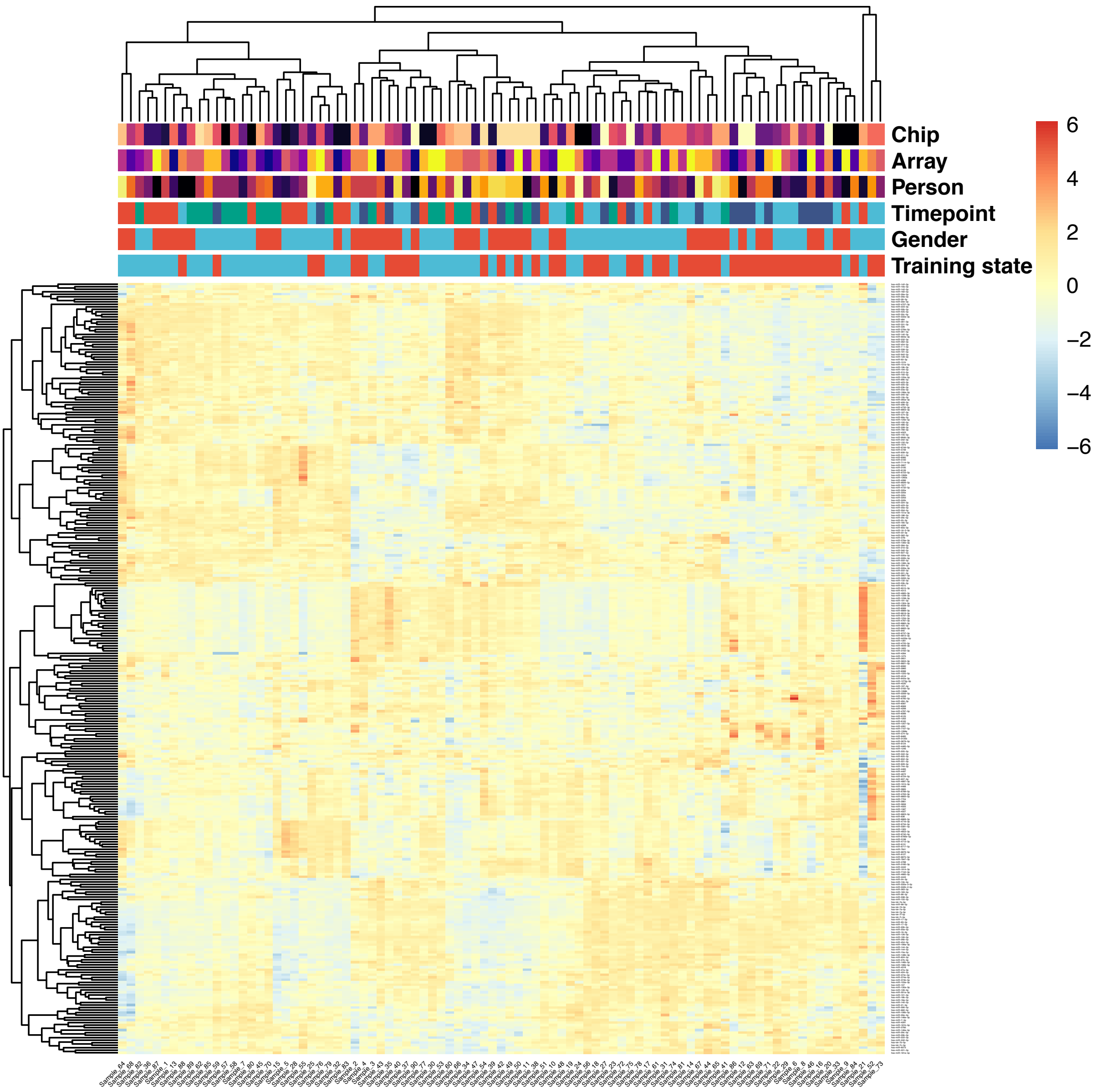


**Figure S2:** MiRNA to sample expression matrix using Z-scores. Higher expression values are color coded in red, while lower expression values are colored in blue. Along each dimension a hierarchical clustering is plotted as a dendrogram, i.e. for samples on top and miRNAs on the left of the heatmap. Color bars on top of the matrix encode for different annotation variables for each of the samples. Several clusters of miRNAs that show a similar expression profile can be identified form the expression matrix and the clustering.

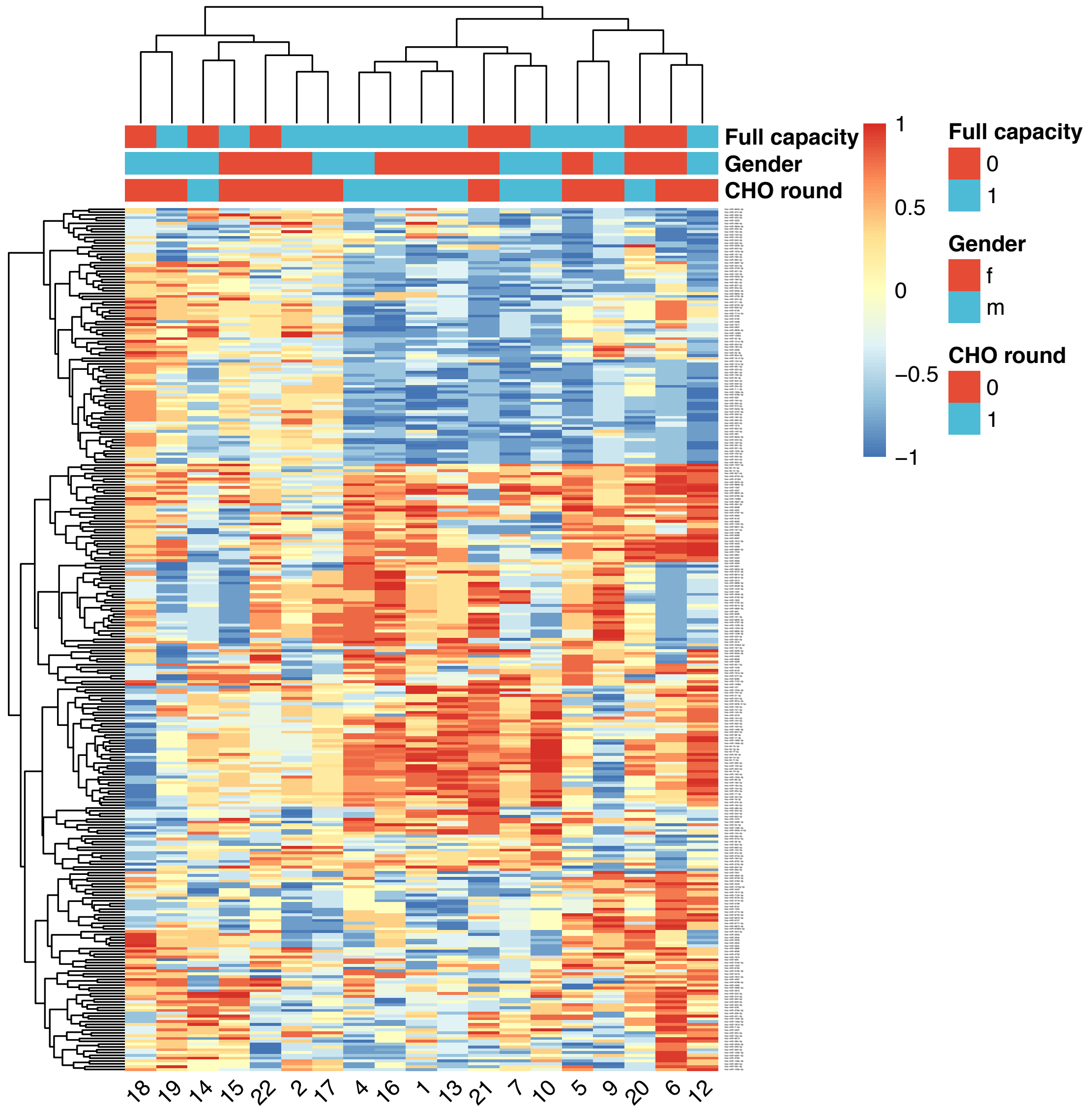


**Figure S3:** Correlation matrix between measured VO2 max levels and miRNA expression distributed over the study participants. A high positive correlation is colored in red, while a strong negative correlation is colored in blue. Along each dimension a hierarchical clustering is plotted as a dendrogram, i.e. for participants on top and miRNAs on the left of the heatmap. Color bars on top of the matrix encode for three binary annotation variables obtained for each of the participants. Similar to Supplementary Figure S2, several clusters of miRNAs that show comparable correlations with VO2 max levels can be identified.

**Table S1**: Blood sample and phenotype parameter measurements from participants across each timepoint of the study.

| Always_full_capacity | CHO_1._Round | Participant | Number of training_sessions_1._Round | Number_of_training_sessions_2._Round | Heart_beat_avg_1._Round |
|---|---|---|---|---|---|
| 1 | 1 | 1 | 30 | 31 | 145 |
| 1 | 1 | 6 | 32 | 32 | 159 |
| 1 | 1 | 2 | 26 | 30 | 159 |
| 1 | 0 | 3 | 31 | 31 | 144 |
| 1 | 0 | 4 | 26 | 26 | 138 |
| 1 | 0 | 7 | 28 | 26 | 142 |
| 0 | 1 | 8 | 30 | 27 | 125 |
| 0 | 0 | 9 | 27 | 27 | 143 |
| 0 | 1 | 10 | 27 | 28 | 144 |
| 0 | 0 | 11 | 28 | 31 | 131 |
| 1 | 0 | 12 | 32 | 35 | 141 |
| 0 | 1 | 13 | 31 | 33 | 152 |
| 0 | 0 | 14 | 27 | 27 | 130 |
| 1 | 1 | 15 | 28 | 28 | 146 |
| 1 | 0 | 16 | 28 | 26 | 155 |
| 0 | 0 | 17 | 32 | 33 | 154 |
| 1 | 1 | 18 | 30 | 32 | 142 |
| 1 | 0 | 19 | 27 | 30 | 152 |
| 1 | 0 | 20 | 28 | 28 | 149 |
| 1 | 0 | 21 | 31 | 30 | 145 |
| 1 | 0 | 22 | 34 | 33 | 146 |
| 1 | 1 | 23 | 32 | 32 | 158 |
| 0 | 1 | 5 | 29 | 29 | 158 |

| Heart_beat_avg_2._Round | Hf-Reserved | E1_Weight_kg | E1_Height_m | E1_BMI | E1_Body_fat |
|---|---|---|---|---|---|
| 140 | 140 | 71 | 1,66 | 25,76571346 | 31,3 |
| 159 | 158 | 75,4 | 1,74 | 24,90421456 | 19,9 |
| 155 | 156 | 92 | 1,74 | 30,3871053 | 25,5 |
| 145 | 148 | 79,1 | 1,645 | 29,23106771 | 24 |
| 139 | 144 | 78,6 | 1,63 | 29,58334902 | 33,2 |
| 145 | 141 | 81,7 | 1,705 | 28,10433347 | 18,6 |
| 126 | 125 | 99,7 | 1,81 | 30,43252648 | 19,9 |
| 145 | 143 | 79 | 1,66 | 28,66889244 | 28 |
| 145 | 144 | 93 | 1,805 | 28,54490067 | 22,3 |
| 134 | 133 | 88,2 | 1,73 | 29,46974506 | 31,4 |
| 145 | 141 | 80 | 1,77 | 25,53544639 | 29,8 |
| 149 | 156 | 77,2 | 1,7 | 26,71280277 | 26,7 |
| 129 | 128 | 90 | 1,8 | 27,77777778 | 21,1 |
| 143 | 145 | 83 | 1,69 | 29,06060712 | 23,3 |
| 154 | 161 | 105,7 | 1,95 | 27,79750164 | 22,4 |
| 151 | 155 | 69,7 | 1,67 | 24,9919323 | 23,5 |
| 141 | 141 | 79,5 | 1,705 | 27,34754603 | 28 |
| 154 | 157 | 72,8 | 1,71 | 24,89654937 | 22,6 |
| 149 | 149 | 92,3 | 1,76 | 29,7972624 | 24,1 |
| 145 | 144 | 72,9 | 1,73 | 24,35764643 | 17,3 |
| 142 | 144 | 82,3 | 1,82 | 24,84603309 | 20,2 |
| 156 | 155 | 81,5 | 1,77 | 26,01423601 | 27,1 |
| 157 | 158 | 97,6 | 1,77 | 31,1532446 | 18,3 |

| E1_VO2max_(ml/min/kg) | E1_VO2max_(l/min.) | E1_VT_l/min | E1_V | E1_Resting-HF | E1_HFmax |
|---|---|---|---|---|---|
| 27,7 | 1,959 | 1,104 | 8,6 | 63 | 180 |
| 51 | 3,831 | 1,713 | 15 | 54 | 200 |
| 40,3 | 3,712 | 2,328 | 13,4 | 75 | 197 |
| 26,7 | 2,112 | 1,246 | 10,2 | 74 | 180 |
| 27 | 2,393 | 1,461 | 10,2 | 56 | 181 |
| 39,7 | 3,192 | 1,706 | 11,8 | 63 | 175 |
| 34,7 | 3,433 | 2,019 | 11 | 56 | 148 |
| 28 | 2,197 | 2,316 | 8,6 | 80 | 170 |
| 34,3 | 3,212 | 1,543 | 10,2 | 59 | 175 |
| 20 | 1,749 | 1,665 | 7,8 | 55 | 166 |
| 28 | 2,231 | 1,003 | 10,2 | 56 | 178 |
| 33 | 2,597 | | 10,2 | 70 | 190 |
| 38,7 | 3,474 | 1,602 | 13,2 | 58 | 165 |
| 31,7 | 2,639 | 1,541 | 11 | 70 | 180 |
| 36,3 | 3,836 | 1,455 | 11,8 | 69 | 200 |
| 33,7 | 2,192 | 1,908 | 9,4 | 73 | 190 |
| 26,7 | 2,109 | 1,62 | 9,4 | 65 | 180 |
| 38,7 | 2,816 | 1,479 | 11,8 | 81 | 190 |
| 32,3 | 2,998 | 1,384 | 11 | 64 | 185 |
| 33 | 2,393 | 1,562 | 11,8 | 72 | 175 |
| 42 | 3,435 | 1,418 | 14,2 | 51 | 184 |
| 37,7 | 3,067 | 1,773 | 12,6 | 69 | 190 |
| 35 | 3,392 | 2,101 | 12,6 | 71 | 190 |

| E1_RQ | E1_bLa_1P | E1_Systolic_blood_pressure_right_auto_1._measurement | E1_Systolic_blood_pressure_right_auto_2._measurement | E1_Avg_systolic_blood_pressure_right_auto | E1_Diastolic_blood_pressure_right_auto_1._measurement |
|---|---|---|---|---|---|
| 1,17 | 8,21 | 148 | 135 | 141,5 | 96 |
| 1,13 | 10,2 | 130 | 121 | 125,5 | 80 |
| 1,29 | 10,8 | 137 | 138 | 137,5 | 78 |
| 1,32 | 6,64 | 142 | 142 | 142 | 100 |
| 1,3 | 9,58 | 108 | 109 | 108,5 | 68 |
| 1,24 | 8,2 | 120 | 120 | 120 | 85 |
| 1,07 | 3,36 | 132 | 126 | 129 | 86 |
| 1,09 | 5,55 | 122 | 132 | 127 | 86 |
| 1,26 | 8,49 | 116 | 112 | 114 | 78 |
| 1,19 | 5,29 | 126 | 118 | 122 | 80 |
| 1,19 | 6,17 | 122 | 125 | 123,5 | 82 |
| 1,11 | 6,55 | 118 | 130 | 124 | 75 |
| 1,16 | 10,7 | 157 | 143 | 150 | 91 |
| 1,25 | 9,29 | 114 | 116 | 115 | 73 |
| 1,26 | 6,94 | 136 | 133 | 134,5 | 90 |
| 1,1 | 7,96 | 122 | 131 | 126,5 | 84 |
| 1,29 | 6,71 | 135 | 134 | 134,5 | 90 |
| 1,24 | 5,61 | 130 | 130 | 130 | 85 |
| 1,26 | 9,05 | 135 | 150 | 142,5 | 90 |
| 1,32 | 6,96 | 134 | 141 | 137,5 | 82 |
| 1,17 | 9,18 | 146 | 140 | 143 | 90 |
| 1,25 | 9,02 | 115 | 118 | 116,5 | 78 |
| 1,29 | 11 | 130 | 122 | 126 | 90 |

| E1_Diastolic_blood_pressure_right_auto_2._measurement | E1_Avg_diastolic_blood_pressure_right_auto | E1_Cholesterol | E1_HDL | E1_LDL | E1_Triglycerides |
|---|---|---|---|---|---|
| 91 | 93,5 | 241 | 63,7 | 145,6 | 182 |
| 85 | 82,5 | 231 | 39,5 | 163 | 107 |
| 84 | 81 | 184 | 28,8 | 117,5 | 204 |
| 102 | 101 | 183 | 53,3 | 113 | 72 |
| 65 | 66,5 | 187 | 34,9 | 144 | 65 |
| 85 | 85 | 230 | 46,7 | 148 | 169 |
| 81 | 83,5 | 250 | 42,1 | 169,9 | 126 |
| 97 | 91,5 | 261 | 51,2 | 182 | 166 |
| 70 | 74 | 207 | 57,7 | 126 | 47 |
| 74 | 77 | 202 | 70,9 | 102 | 78 |
| 82 | 82 | 222 | 95,7 | 103 | 66 |
| 82 | 78,5 | 156 | 47,1 | 98,1 | 53 |
| 91 | 91 | 233 | 39,3 | 173 | 96 |
| 78 | 75,5 | 207 | 41,5 | 130 | 143 |
| 91 | 90,5 | 222 | 41 | 160 | 61 |
| 89 | 86,5 | 238 | 75,2 | 136 | 47 |
| 94 | 92 | 268 | 62,3 | 192 | 102 |
| 86 | 85,5 | 259 | 54,2 | 180 | 145 |
| 97 | 93,5 | 273 | 40,6 | 183,1 | 214 |
| 85 | 83,5 | 202 | 32,4 | 143 | 153 |
| 98 | 94 | 185 | 57,9 | 109 | 38 |
| 82 | 80 | 166 | 49 | 103 | 45 |
| 86 | 88 | 166 | 25,8 | 110 | 184 |

| E1_Glucose | E1_uric_acid | E1_Ferritin | E1_GOT | E1_GPT | E1_Gamma-GT |
|---|---|---|---|---|---|
| 79 | 4,1 | 19,4 | 9 | 12 | 8 |
| 91 | 4,5 | 62,9 | 11 | 19 | 19 |
| 95 | 7,9 | 124 | 16 | 34 | 36 |
| 97 | 4,2 | 102 | 18 | 32 | 19 |
| 88 | 5,5 | 27,5 | 21 | 27 | 21 |
| 99 | 6,4 | 313 | 32 | 57 | 29 |
| 101 | 6,9 | 269,2 | 14 | 26 | 26 |
| 104 | 6 | 86,2 | 14 | 17 | 26 |
| 106 | 6,5 | 197 | 23 | 65 | 137 |
| 87 | 3,5 | 23,7 | 16 | 22 | 21 |
| 96 | 4,9 | 13,4 | 9 | 19 | 17 |
| 82 | 4,2 | 39,9 | 15 | 23 | 21 |
| 100 | 6,3 | 151 | 15 | 34 | 76 |
| 92 | 6,1 | 173 | 28 | 55 | 55 |
| 87 | 5,6 | 164 | 16 | 28 | 31 |
| 89 | 4,9 | 10,7 | 12 | 16 | 15 |
| 87 | 4,7 | 44,4 | 13 | 28 | 51 |
| 98 | 5,6 | 220 | 15 | 36 | 36 |
| 76 | 5,5 | 215,1 | 19 | 36 | 87 |
| 107 | 7,1 | 246 | 13 | 25 | 20 |
| 97 | 4,9 | 107 | 19 | 33 | 47 |
| 91 | 4,8 | 26,9 | 11 | 16 | 9 |
| 103 | 6,3 | 270 | 29 | 36 | 21 |

| E1_urea | E1_creatinine | E1_BSG | E1_Hb | E1_HKT | E1_60P_Cholesterol |
|---|---|---|---|---|---|
| 13 | 0,56 | 13 | 13,4 | 41,7 | 240 |
| 34 | 0,87 | 4 | 13,1 | 39,2 | 224 |
| 29 | 0,88 | 6 | 13,9 | 42,5 | 180 |
| 21 | 0,79 | 3 | 14,3 | 43,9 | 184 |
| 29 | 0,74 | 15 | 12,4 | 39,8 | 188 |
| 43 | 1,14 | 1 | 16,1 | 47,9 | 220 |
| 52 | 1,08 | 1 | 14,5 | 46 | 255 |
| 23 | 0,65 | 6 | 13,9 | 42,5 | 259 |
| 33 | 0,82 | 3 | 14,3 | 43,6 | 204 |
| 22 | 0,7 | 19 | 12,1 | 37 | 193 |
| 27 | 0,68 | 5 | 12,5 | 38,9 | 195 |
| 27 | 0,67 | 7 | 122 | 38,8 | 152 |
| 36 | 1,02 | 3 | 14,6 | 42,6 | 217 |
| 32 | 1,01 | 2 | 14,4 | 43,7 | 203 |
| 26 | 0,71 | 5 | 14,7 | 43,7 | 220 |
| 33 | 0,63 | 9 | 13,9 | 42,4 | 195 |
| 27 | 0,71 | 6 | 13,1 | 40,7 | 270 |
| 33 | 0,89 | 2 | 13,8 | 42,2 | 257 |
| 30 | 0,86 | 3 | 14,2 | 43,7 | 273 |
| 37 | 0,65 | 2 | 14,7 | 44,8 | 217 |
| 35 | 0,72 | 1 | 16,2 | 49,4 | 183 |
| 19 | 0,71 | 2 | 13,1 | 40,2 | 164 |
| 30 | 1,16 | 7 | 15,1 | 44,9 | 167 |

| E1_60P_HDL | E1_60P_LDL | E1_60P_Triglycerides | E1_60P_Glucose | E1_60P_BSG | E1_60P_Hb |
|---|---|---|---|---|---|
| 63,7 | 145,6 | 162 | 84 | 7 | 13,4 |
| 38,9 | 155 | 95 | 84 | 4 | 12,8 |
| 28,8 | 116,2 | 218 | 86 | 5 | 14,1 |
| 53 | 112 | 75 | 103 | 3 | 14,5 |
| 33,8 | 143 | 64 | 83 | 15 | 13 |
| 46,2 | 146 | 133 | 86 | 1 | 15,5 |
| 44,9 | 173,3 | 91 | 89 | 1 | 14,5 |
| 52,3 | 183 | 141 | 96 | 10 | 13,8 |
| 58,7 | 125 | 39 | 99 | 3 | 14,1 |
| 66,3 | 102 | 77 | 80 | 18 | 11,9 |
| 85,8 | 91,4 | 51 | 89 | 6 | 12,5 |
| 46,3 | 98,4 | 49 | 87 | 9 | 12,3 |
| 35,2 | 161 | 93 | 86 | | 14,3 |
| 40,2 | 127 | 132 | 88 | 4 | 14,4 |
| 41,3 | 161 | 59 | 80 | 5 | 14,9 |
| 63,1 | 110 | 33 | 76 | 7 | 12,8 |
| 64,8 | 194 | 98 | 83 | 12 | 13,3 |
| 54,2 | 176 | 143 | 89 | 3 | 13,8 |
| 40,6 | 183,1 | 208 | 76 | 5 | 14,2 |
| 36 | 150 | 171 | 100 | 2 | 15,2 |
| 54,7 | 103 | 43 | 85 | 1 | 16 |
| 47,5 | 100 | 32 | 84 | 3 | 12,9 |
| 27,5 | 109 | 192 | 89 | 8 | 15,1 |

| E1_60P_HKT | E1_60_P_Systolic_blood_pressure_right_auto_1._measurement | E1_60_P_Systolic_blood_pressure_right_auto_2._measurement | E1_60_P_Avg_systolic_blood_pressure_right_auto | E1_60_P_Diastolic_blood_pressure_right_auto_1._measurement | E1_60_P_Diastolic_blood_pressure_right_auto_2._measurement |
|---|---|---|---|---|---|
| 41,7 | 144 | 134 | 139 | 98 | 98 |
| 38,2 | 117 | 121 | 119 | 66 | 66 |
| 42 | 133 | 139 | 136 | 78 | 85 |
| 45,2 | 123 | 119 | 121 | 86 | 90 |
| 40,8 | 108 | 108 | 108 | 74 | 62 |
| 45,9 | 138 | 126 | 132 | 90 | 82 |
| 46 | 143 | 140 | 141,5 | 89 | 94 |
| 42,3 | 130 | 125 | 127,5 | 91 | 101 |
| 42,7 | 115 | 120 | 117,5 | 71 | 78 |
| 36,3 | 111 | 114 | 112,5 | 75 | 70 |
| 38,2 | 114 | 118 | 116 | 80 | 80 |
| 38,5 | 113 | 118 | 115,5 | 77 | 79 |
| 43 | 124 | 122 | 123 | 76 | 82 |
| 44 | 110 | 116 | 113 | 71 | 80 |
| 44,9 | 122 | 122 | 122 | 77 | 76 |
| 39 | 109 | 118 | 113,5 | 75 | 78 |
| 41,7 | 138 | 136 | 137 | 94 | 92 |
| 42,1 | 122 | 122 | 122 | 78 | 78 |
| 43,7 | 131 | 138 | 134,5 | 94 | 89 |
| 45,7 | 130 | 131 | 130,5 | 89 | 90 |
| 48,5 | 118 | 167 | 142,5 | 80 | 86 |
| 39,9 | 124 | 110 | 117 | 78 | 75 |
| 45,2 | 123 | 126 | 124,5 | 89 | 86 |

| E1_60_P_Avg_diastolic_blood_pressure_right_auto | A1_Weight_kg | A1_Height_m | A1_BMI | A1_Body_fat | A1_VO2max_(ml/min/kg) |
|---|---|---|---|---|---|
| 98 | 70,2 | 1,66 | 25,47539556 | 29,4 | 28,7 |
| 66 | 70,9 | 1,74 | 23,41788876 | 17,6 | 46,3 |
| 81,5 | 93,3 | 1,74 | 30,81648831 | 26,3 | 41,7 |
| 88 | 81,1 | 1,66 | 29,43097692 | 25,4 | 33,3 |
| 68 | 78,1 | 1,625 | 29,57633136 | 31,4 | 28 |
| 86 | 80,6 | 1,73 | 26,93040195 | 20,7 | 44 |
| 91,5 | 97,8 | 1,795 | 30,35358199 | 23 | 38,7 |
| 96 | 79,8 | 1,67 | 28,61343182 | 27,5 | 28 |
| 74,5 | 94,6 | 1,8 | 29,19753086 | 23,6 | 35 |
| 72,5 | 86,7 | 1,75 | 28,31020408 | 27,5 | 25,6 |
| 80 | 79 | 1,78 | 24,93372049 | 28,5 | 34,6 |
| 78 | 77,3 | 1,7 | 26,74740484 | 25,7 | 35,3 |
| 79 | 92,7 | 1,79 | 28,93168128 | 20,7 | 38,7 |
| 75,5 | 80,2 | 1,68 | 28,41553288 | 21 | 34,7 |
| 76,5 | 103,9 | 1,96 | 27,04602249 | 20,9 | 39,3 |
| 76,5 | 67,2 | 1,675 | 23,95188238 | 26 | 33 |
| 93 | 80,8 | 1,695 | 28,12366756 | 28 | 32 |
| 78 | 70,5 | 1,72 | 23,83044889 | 20,1 | 45 |
| 91,5 | 91,1 | 1,745 | 29,91765256 | 22,6 | 34,7 |
| 89,5 | 73,7 | 1,735 | 24,48321969 | 16,5 | 33,3 |
| 83 | 76,6 | 1,81 | 23,38145966 | 17,4 | 43 |
| 76,5 | 81,8 | 1,77 | 26,10999394 | 26 | 39,3 |
| 87,5 | 95,8 | 1,77 | 30,57869705 | 18 | 43,3 |

| A1_VO2max_(l /min.) | A1_VT_l/min | A1_V | A1_Resting-HF | A1_HFmax | A1_RQ |
|---|---|---|---|---|---|
| 2,035 | 1,085 | 9,4 | 59 | 175 | 1,12 |
| 3,495 | 2,111 | 15 | 50 | 190 | 1,16 |
| 3,777 | 2,553 | 12,6 | 77 | 196 | 1,21 |
| 2,707 | 1,487 | 11 | 68 | 185 | 1,21 |
| 2,173 | 1,494 | 11 | 64 | 185 | 1,31 |
| 3,593 | 2,577 | 13,4 | 58 | 175 | 1,21 |
| 3,846 | 2,146 | 11,8 | 57 | 160 | 1,13 |
| 2,234 | 1,884 | 9,4 | 79 | 170 | 1,06 |
| 3,302 | 1,593 | 11 | 54 | 170 | 1,2 |
| 2,232 | 2,025 | 9,4 | 60 | 165 | 1,1 |
| 2,758 | 1,301 | 11,8 | 54 | 160 | 1,1 |
| 2,787 |  | 10,2 | 57 | 185 | 1,13 |
| 3,565 | 1,774 | 13,2 | 60 | 155 | 1,17 |
| 2,873 | 1,85 | 11,8 | 58 | 161 | 1,15 |
| 4,041 | 1,563 | 12,6 | 54 | 184 | 1,24 |
| 2,295 | 2,007 | 10,2 | 69 | 191 | 1,2 |
| 2,561 | 1,671 | 10,2 | 57 | 180 | 1,29 |
| 3,29 | 1,456 | 13,4 | 63 | 172 | 1,1 |
| 3,149 | 2,255 | 11,8 | 60 | 180 | 1,24 |
| 2,446 | 1,721 | 12,6 | 64 | 170 | 1,31 |
| 3,29 | 1,573 | 15,8 | 45 | 172 | 1,29 |
| 3,217 | 2,038 | 13,4 | 65 | 195 | 1,15 |
| 3,751 | 2,663 | 14,2 | 67 | 170 | 1,19 |

| A1_bLa_1P | A1_Systolic_blood_pressure_right_auto_1._measurement | A1_Systolic_blood_pressure_right_auto_2._measurement | A1_Avg_systolic_blood_pressure_right_auto | A1_Diastolic_blood_pressure_right_auto_1._measurement | A1_Diastolic_blood_pressure_right_auto_2._measurement |
|---|---|---|---|---|---|
| 9,43 | 125 | 125 | 125 | 86 | 85 |
| 8,06 | 126 | 128 | 127 | 87 | 79 |
| 10,9 | 131 | 133 | 132 | 73 | 72 |
| 8,26 | 134 | 143 | 138,5 | 96 | 101 |
| 7,93 | 110 | 110 | 110 | 70 | 68 |
| 8,2 | 123 | 134 | 128,5 | 85 | 82 |
| 5,21 | 125 | 134 | 129,5 | 85 | 80 |
| 5,35 | 122 | 118 | 120 | 74 | 75 |
| 7,21 | 120 | 115 | 117,5 | 80 | 72 |
| 7,04 | 126 | 119 | 122,5 | 96 | 82 |
| 6,88 | 124 | 132 | 128 | 78 | 83 |
| 6,62 | 122 | 122 | 122 | 78 | 82 |
| 7,16 | 142 | 138 | 140 | 90 | 82 |
| 5,89 | 117 | 119 | 118 | 76 | 76 |
| 8,59 | 133 | 133 | 133 | 79 | 83 |
| 6,9 | 111 | 121 | 116 | 82 | 82 |
| 7,39 | 136 | 134 | 135 | 95 | 99 |
| 5,94 | 123 | 120 | 121,5 | 70 | 71 |
| 10,4 | 138 | 150 | 144 | 94 | 93 |
| 10,9 | 127 | 130 | 128,5 | 81 | 81 |
| 9,32 | 132 | 130 | 131 | 84 | 82 |
| 8,01 | 121 | 119 | 120 | 84 | 86 |
| 6,54 | 134 | 134 | 134 | 93 | 94 |

| A1_Avg_diastolic_blood_pressure_right_auto | A1_Cholesterol | A1_HDL | A1_LDL | A1_Triglycerides | A1_Glucose |
|---|---|---|---|---|---|
| 85,5 | 236 | 61,5 | 148 | 205 | 92 |
| 83 | 184 | 39,9 | 120,9 | 64 | 93 |
| 72,5 | 189 | 34,9 | 125 | 143 | 99 |
| 98,5 | NA | NA | NA | NA | NA |
| 69 | 222 | 40,6 | 162 | 87 | 89 |
| 83,5 | 235 | 52 | 167 | 126 | 99 |
| 82,5 | 212 | 45,5 | 136 | 102 | 102 |
| 74,5 | 292 | 59 | 195 | 193 | 97 |
| 76 | 196 | 59,1 | 125 | 41 | 107 |
| 89 | 253 | 86,9 | 138 | 37 | 102 |
| 80,5 | 220 | 102 | 99,6 | 46 | 96 |
| 80 | 151 | 34,2 | 103 | 52 | 91 |
| 86 | 222 | 35,2 | 145 | 182 | 102 |
| 76 | 212 | 54,7 | 130 | 114 | 90 |
| 81 | 195 | 37,9 | 143 | 65 | 96 |
| 82 | 213 | 76 | 112 | 55 | 77 |
| 97 | 240 | 76,4 | 157 | 87 | 86 |
| 70,5 | 227 | 55,5 | 148 | 137 | 106 |
| 93,5 | 299 | 44,9 | 177 | 353 | 87 |
| 81 | 217 | 34,8 | 157 | 127 | 102 |
| 83 | 136 | 55,5 | 60,9 | 21 | 94 |
| 85 | 172 | 53,1 | 97,1 | 67 | 92 |
| 93,5 | 186 | 28,4 | 121 | 242 | 96 |

| A1_uric_acid | A1_Ferritin | A1_GOT | A1_GPT | A1_Gamma-GT | A1_urea |
|---|---|---|---|---|---|
| 4,3 | 19,3 | 10 | 13 | 9 | 18 |
| 4,8 | 72,4 | 20 | 24 | 23 | 23 |
| 7,6 | 101 | 14 | 34 | 37 | 39 |
| NA | NA | NA | NA | NA | NA |
| 5,3 | 33,9 | 31 | 49 | 32 | 24 |
| 5,7 | 240 | 21 | 38 | 21 | 34 |
| 7 | 238 | 14 | 22 | 21 | 48 |
| 5,8 | 62,2 | 16 | 21 | 25 | 24 |
| 6 | 213 | 32 | 84 | 166 | 41 |
| 5 | 24,5 | 14 | 17 | 24 | 34 |
| 4,9 | 7,2 | 13 | 25 | 17 | 32 |
| 4,8 | 13,1 | 26 | 38 | 21 | 27 |
| 6,9 | 97,6 | 24 | 56 | 247 | 37 |
| 5,7 | 140 | 21 | 44 | 39 | 31 |
| 6,3 | 102 | 10 | 23 | 22 | 21 |
| 4,8 | 8,7 | 15 | 17 | 14 | 33 |
| 5,2 | 18,8 | 14 | 18 | 25 | 21 |
| 5,6 | 161 | 13 | 21 | 34 | 32 |
| 5 | 154 | 17 | 34 | 58 | 26 |
| 7,1 | 191 | 24 | 29 | 26 | 42 |
| 4,5 | 41 | 12 | 24 | 30 | 36 |
| 4 | 17,1 | 10 | 17 | 11 | 18 |
| 6,6 | 167 | 38 | 38 | 20 | 51 |

| A1_creatinine | A1_BSG | A1_Hb | A1_HKT | E2_Weight_kg | E2_Height_m |
|---|---|---|---|---|---|
| 0,62 | 17 | 12,4 | 37,5 | 70,3 | 1,665 |
| 0,93 | 4 | 12,7 | 37,3 | 80 | 1,725 |
| 0,78 | 4 | 14,4 | 40,6 | 93,5 | 1,74 |
| NA | NA | NA | NA | 81,6 | 1,66 |
| 0,66 | 20 | 13,4 | 41,6 | 72,1 | 1,63 |
| 1,08 | 1 | 15,6 | 47,9 | 80,3 | 1,71 |
| 1,03 | 1 | 14,9 | 45,9 | 96 | 1,81 |
| 0,72 | 5 | 13,9 | 41,6 | 78,9 | 1,66 |
| 0,8 | 3 | 14 | 41,7 | 91,6 | 1,78 |
| 0,71 | 10 | 12,5 | 37,3 | 81,4 | 1,76 |
| 0,61 | 5 | 12,7 | 38,7 | 82,9 | 1,78 |
| 0,78 | 11 | 12,3 | 35,2 | 77,9 | 1,69 |
| 1 | 2 | 14,3 | 42,6 | 85,3 | 1,79 |
| 0,98 | 2 | 14,4 | 45,7 | 82,7 | 1,7 |
| 0,69 | 4 | 13,8 | 42,4 | 104,8 | 1,95 |
| 0,63 | 8 | 13,1 | 40,2 | 64,5 | 1,675 |
| 0,73 | 5 | 12 | 39,1 | 78,2 | 1,7 |
| 0,88 | 2 | 13,3 | 41,7 | 69,3 | 1,72 |
| 0,88 | 2 | 145 | 43,7 | 92,5 | 1,755 |
| 0,81 | 2 | 14,5 | 45,8 | 74,1 | 1,73 |
| 0,89 | 2 | 14,3 | 42,4 | 80 | 1,82 |
| 0,59 | 4 | 13 | 37,1 | 81,3 | 1,775 |
| 1 | 11 | 15,2 | 43,8 | 99,5 | 1,77 |

| E2_BMI | E2_Body_fat | E2_VO2max_(ml/min/kg) | E2_VO2max_(l/min.) | E2_VT_l/min | E2_V |
|---|---|---|---|---|---|
| 25,35869203 | 28,25 | 29 | 2,073 | 1,127 | 8,6 |
| 26,88510817 | 20,4 | 46,3 | 3,702 | 2,077 | 12,6 |
| 30,88254723 | 26,8 | 37 | 3,503 | 2,217 | 11,8 |
| 29,61242561 | 25,4 | 33 | 2,685 | 1,246 | 11 |
| 27,13688886 | 31 | 34 | 2,447 | 1,631 | 11 |
| 27,46144113 | 20,1 | 41,3 | 3,305 | 1,521 | 13,4 |
| 29,30313482 | 17,45 | 37 | 3,593 | 2,441 | 11 |
| 28,6326027 | 26 | 24,3 | 1,915 | 1,917 | 9,4 |
| 28,9104911 | 22,6 | 29,7 | 2,706 | 1,383 | 10,2 |
| 26,27840909 | 24,1 | 25 | 2,183 | 1,395 | 10,2 |
| 26,16462568 | 30,4 | 29,7 | 2,454 | 1,106 | 11 |
| 27,27495536 | 23,7 | 28,7 | 2,23 | | 8,6 |
| 26,62214038 | 17,2 | 36,3 | 3,348 | 1,606 | 14,2 |
| 28,61591696 | 22,2 | 30,3 | 2,5 | 1,64 | 11 |
| 27,56081525 | 20,1 | 37 | 3,859 | 1,049 | 12,6 |
| 22,98952996 | 19,1 | 36,3 | 2,341 | 2,275 | 11 |
| 27,05882353 | 26,5 | 25,7 | 2,013 | 1,64 | 9,4 |
| 23,42482423 | 19,4 | 38 | 2,747 | 1,504 | 13,4 |
| 30,03222376 | 24,1 | 32,7 | 3,018 | 1,6 | 11,8 |
| 24,75859534 | 17,4 | 34 | 2,502 | 1,788 | 11,8 |
| 24,1516725 | 18,7 | 50,7 | 3,885 | 1,399 | 15 |
| 25,80440389 | 24,8 | 38,7 | 3,142 | 1,476 | 12,6 |
| 31,75971145 | 23,5 | 35,7 | 3,544 | 2,018 | 12,6 |

| E2_Resting-HF | E2_HFmax | E2_RQ | E2_bLa_1P | E2_Systolic_blood_pressure_right_auto_1._measurement | E2_Systolic_blood_pressure_right_auto_2._measurement |
|---|---|---|---|---|---|
| 66 | 172 | 1,13 | 8,2 | 138 | 127 |
| 61 | 200 | 1,16 | 8,23 | 136 | 140 |
| 78 | 190 | 1,17 | 10,4 | 130 | 124 |
| 68 | 191 | 1,32 | 7,99 | 142 | 148 |
| 55 | 185 | 1,36 | 7,71 | 112 | 114 |
| 55 | 172 | 1,26 | 9,61 | 131 | 120 |
| 49 | 158 | 1,04 | 5,71 | 127 | 134 |
| 75 | 176 | 1,17 | 6,75 | 122 | 120 |
| 59 | 180 | 1,27 | 8,09 | 123 | 122 |
| 64 | 185 | 1,25 | 5,78 | 133 | 130 |
| 64 | 180 | 1,21 | 6,54 | 128 | 138 |
| 65 | 195 | 1,21 | 4,89 | 113 | 114 |
| 54 | 160 | 1,22 | 7,3 | 122 | 131 |
| 63 | 180 | 1,32 | 6,48 | 132 | 110 |
| 56 | 185 | 1,2 | 8,17 | 127 | 126 |
| 57 | 188 | 1,2 | 6,09 | 129 | 124 |
| 56 | 178 | 1,31 | 7,22 | 146 | 142 |
| 56 | 185 | 1,21 | 9,39 | 123 | 121 |
| 61 | 183 | 1,23 | 8,87 | 142 | 134 |
| 64 | 180 | 1,28 | 8,11 | 130 | 130 |
| 44 | 175 | 1,14 | 7,5 | 139 | 134 |
| 72 | 190 | 1,19 | 11,1 | 133 | 129 |
| 83 | 190 | 1,37 | 8,65 | 138 | 138 |

| E2_Avg_systolic_blood_pressure_right_auto | E2_Diastolic_blood_pressure_right_auto_1._measurement | E2_Diastolic_blood_pressure_right_auto_2._measurement | E2_Avg_diastolic_blood_pressure_right_auto | E2_Cholesterol | E2_HDL |
|---|---|---|---|---|---|
| 132,5 | 97 | 92 | 94,5 | 253 | 58,6 |
| 138 | 92 | 87 | 89,5 | 250 | 40,4 |
| 127 | 74 | 74 | 74 | 230 | 34,5 |
| 145 | 97 | 102 | 99,5 | 246 | 62,2 |
| 113 | 78 | 72 | 75 | 181 | 37,8 |
| 125,5 | 85 | 78 | 81,5 | 197 | 47,5 |
| 130,5 | 82 | 89 | 85,5 | 233 | 48,9 |
| 121 | 84 | 79 | 81,5 | 246 | 48,4 |
| 122,5 | 79 | 78 | 78,5 | 154 | 38,3 |
| 131,5 | 90 | 97 | 93,5 | 250 | 98,1 |
| 133 | 87 | 82 | 84,5 | 223 | 97,2 |
| 113,5 | 77 | 74 | 75,5 | 145 | 25,8 |
| 126,5 | 76 | 88 | 82 | 199 | 33,8 |
| 121 | 88 | 80 | 84 | 219 | 46,1 |
| 126,5 | 82 | 76 | 79 | 208 | 46 |
| 126,5 | 81 | 80 | 80,5 | 174 | 63,9 |
| 144 | 100 | 101 | 100,5 | 250 | 73,3 |
| 122 | 74 | 70 | 72 | 258 | 51,3 |
| 138 | 83 | 82 | 82,5 | 309 | 47,1 |
| 130 | 84 | 82 | 83 | 197 | 33,6 |
| 136,5 | 86 | 93 | 89,5 | 185 | 70,5 |
| 131 | 98 | 89 | 93,5 | 180 | 52,3 |
| 138 | 97 | 92 | 94,5 | 215 | 28,3 |

| E2_LDL | E2_Triglycerides | E2_Glucose | E2_uric_acid | E2_Ferritin | E2_GOT |
|---|---|---|---|---|---|
| 162 | 228 | 94 | 4,3 | 24,9 | 8 |
| 169 | 135 | 92 | 4,7 | 103 | 12 |
| 143 | 247 | 98 | 8,3 | 140 | 20 |
| 161 | 110 | 109 | 4,5 | 80,2 | 16 |
| 132 | 31 | 95 | 5,3 | 43,2 | 28 |
| 123 | 181 | 93 | 4,8 | 198 | 24 |
| 162 | 102 | 92 | 6,6 | 240 | 13 |
| 169,1 | 180 | 94 | 6,3 | 75,2 | 14 |
| 101 | 31 | 97 | 6,5 | 324 | 20 |
| 131,6 | 42 | 78 | 3,5 | 19,3 | 12 |
| 106,8 | 64 | 95 | 5 | 6,8 | 16 |
| 103 | 87 | 85 | 4,1 | 28,8 | 50 |
| 136 | 121 | 101 | 5,9 | 95,3 | 16 |
| 122 | 200 | 92 | 5,4 | 207 | 26 |
| 154 | 45 | 81 | 6,2 | 123 | 12 |
| 89,5 | 46 | 89 | 4,3 | 7,9 | 13 |
| 156 | 56 | 82 | 4,9 | 21,4 | 13 |
| 173 | 182 | 97 | 5,3 | 200 | 13 |
| 212 | 361 | 99 | 5,4 | 185 | 21 |
| 126 | 207 | 101 | 7,2 | 215 | 21 |
| 100 | 39 | 93 | 4,3 | 106 | 12 |
| 108 | 83 | 97 | 4 | 22,2 | 5 |
| 141 | 225 | 106 | 7 | 375 | 37 |

| E2_GPT | E2_Gamma-GT | E2_urea | E2_creatinine | E2_BSG | E2_Hb |
|---|---|---|---|---|---|
| 10 | 8 | 16 | 0,73 | 27 | 13,4 |
| 18 | 25 | 28 | 0,95 | 5 | 13,8 |
| 37 | 48 | 30 | 0,86 | 5 | 13,6 |
| 24 | 16 | 26 | 0,77 | 2 | 14,1 |
| 43 | 20 | 31 | 0,5 | 17 | 13,2 |
| 37 | 20 | 40 | 1,23 | 2 | 14,1 |
| 24 | 24 | 42 | 0,94 | 2 | 15,6 |
| 19 | 31 | 34 | 0,73 | 9 | 14,3 |
| 52 | 79 | 29 | 0,88 | 5 | 14 |
| 15 | 19 | 39 | 0,65 | 10 | 13,5 |
| 27 | 16 | 28 | 0,56 | 7 | 12,5 |
| 95 | 24 | 31 | 0,59 | 19 | 13,3 |
| 30 | 66 | 45 | 0,96 | 2 | 13,9 |
| 52 | 53 | 34 | 0,88 | 2 | 15,7 |
| 24 | 31 | 26 | 0,72 | 2 | 14,5 |
| 17 | 15 | 28 | 0,56 | 6 | 12,5 |
| 18 | 33 | 20 | 0,73 | 4 | 12,7 |
| 20 | 28 | 31 | 0,84 | 3 | 14,5 |
| 35 | 63 | 25 | 0,89 | 2 | 14,8 |
| 34 | 19 | 38 | 0,83 | 2 | 14,4 |
| 27 | 37 | 27 | 0,71 | 2 | 16,2 |
| 14 | 11 | 16 | 0,62 | 2 | 13,3 |
| 69 | 41 | 39 | 1,12 | 10 | 15,9 |

| E2_HKT | E2_60P_Cholesterol | E2_60P_HDL | E2_60P_LDL | E2_60P_Triglycerides | E2_60P_Glucose |
|---|---|---|---|---|---|
| 40,7 | 255 | 59,8 | 170 | 213 | 93 |
| 41,8 | 240 | 38,2 | 162 | 146 | 85 |
| 41,7 | 234 | 33,2 | 154 | 243 | 92 |
| 44,7 | 253 | 63,6 | 168 | 112 | 117 |
| 40,6 | 173 | 34,9 | 124 | 31 | 82 |
| 44,5 | 186 | 46,1 | 122 | 119 | 84 |
| 47,4 | 215 | 43,6 | 149 | 77 | 89 |
| 43 | 250 | 50,8 | 174,1 | 168 | 84 |
| 43,1 | 155 | 40,3 | 100 | 34 | 86 |
| 41,7 | 241 | 94,7 | 128,3 | 38 | 78 |
| 37 | 223 | 101,7 | 108,3 | 52 | 83 |
| 39,1 | 146 | 27 | 105 | 79 | 90 |
| 41,9 | 197 | 34,3 | 137 | 85 | 95 |
| 47 | 240 | 43,6 | 123 | 176 | 80 |
| 41,1 | 219 | 49,6 | 159 | 49 | 77 |
| 38,4 | 170 | 62,8 | 86,6 | 46 | 79 |
| 39,9 | 272 | 83,6 | 171 | 62 | 81 |
| 44,2 | 238 | 47,4 | 159 | 139 | 89 |
| 45,1 | 300 | 45,5 | 205 | 337 | 96 |
| 43,9 | 197 | 31,7 | 129 | 217 | 101 |
| 49,2 | 183 | 71,4 | 101 | 32 | 82 |
| 40 | 182 | 55,6 | 110 | 70 | 87 |
| 47,4 | 221 | 28,3 | 144 | 229 | 108 |

| E2_60P_BSG | E2_60P_Hb | E2_60P_HKT | E2_60_P_Systolic_blood_pressure_right_auto_1._measurement | E2_60_P_Systolic_blood_pressure_right_auto_2._measurement | E2_60_P_Avg_systolic_blood_pressure_right_auto |
|---|---|---|---|---|---|
| 28 | 13,7 | 42 | 123 | 130 | 126,5 |
| 5 | 13,8 | 41,8 | 133 | 119 | 126 |
| 6 | 13,7 | 41,7 | 121 | 129 | 125 |
| | 14,4 | 46,5 | 177 | 118 | 147,5 |
| 12 | 12,7 | 39,6 | 114 | 106 | 110 |
| 1 | 14,4 | 45,3 | 130 | 122 | 126 |
| 3 | 14,4 | 44,3 | 140 | 132 | 136 |
| 10 | 14,4 | 42,6 | 118 | 135 | 126,5 |
| 5 | 14,4 | 42,6 | 116 | 116 | 116 |
| 16 | 13,4 | 41,7 | 123 | 131 | 127 |
| 7 | 12,7 | 37,2 | 130 | 119 | 124,5 |
| | 13,2 | 38,3 | 112 | 114 | 113 |
| 3 | 13,7 | 41,4 | 116 | 114 | 115 |
| | 15,9 | 47,4 | 113 | 118 | 115,5 |
| 2 | 14,5 | 41,1 | 134 | 118 | 126 |
| 6 | 12,7 | 39,2 | 106 | 114 | 110 |
| 5 | 13,8 | 43 | 156 | 150 | 153 |
| 3 | 13,9 | 42,3 | 121 | 122 | 121,5 |
| 1 | 14,7 | 45,3 | 135 | 130 | 132,5 |
| 2 | 14,5 | 44,5 | 113 | 115 | 114 |
| 1 | 16 | 48,3 | 130 | 129 | 129,5 |
| 3 | 13,2 | 39,8 | 117 | 112 | 114,5 |
| 10 | 16,2 | 47,4 | 119 | 113 | 116 |

| E2_60_P_Diastolic_blood_pressure_right_auto_1._measurement | E2_60_P_Diastolic_blood_pressure_right_auto_2._measurement | E2_60P_Avg_diastolic_blood_pressure_right_auto | A2_Weight_kg | A2_Height_m | A2_BMI |
|---|---|---|---|---|---|
| 93 | 97 | 95 | 70,9 | 1,67 | 25,42220947 |
| 89 | 77 | 83 | 78,4 | 1,735 | 26,04456478 |
| 78 | 82 | 80 | 94,2 | 1,75 | 30,75918367 |
| 84 | 87 | 85,5 | 81,7 | 1,66 | 29,64871534 |
| 82 | 72 | 77 | 67,9 | 1,625 | 25,71360947 |
| 79 | 78 | 78,5 | 1,72 | 1,79 | 0,536812209 |
| 86 | 82 | 84 | 96,4 | 1,81 | 29,42523122 |
| 82 | 90 | 86 | 78,5 | 1,65 | 28,83379247 |
| 74 | 76 | 75 | 90,8 | 1,79 | 28,33869105 |
| 89 | 91 | 90 | 86,3 | 1,75 | 28,17959184 |
| 85 | 82 | 83,5 | 80,6 | 1,78 | 25,43870723 |
| 81 | 81 | 81 | 79 | 1,7 | 27,33564014 |
| 86 | 70 | 78 | 86,7 | 1,79 | 27,05908055 |
| 78 | 80 | 79 | 82,8 | 1,69 | 28,99058156 |
| 82 | 76 | 79 | 103,8 | 1,95 | 27,29783037 |
| 78 | 75 | 76,5 | 62,7 | 1,69 | 21,95301285 |
| 105 | 105 | 105 | 78,6 | 1,7 | 27,19723183 |
| 76 | 74 | 75 | 69,8 | 1,72 | 23,59383451 |
| 83 | 83 | 83 | 93,1 | 1,755 | 30,22702738 |
| 77 | 78 | 77,5 | 73,2 | 1,73 | 24,45788366 |
| 76 | 89 | 82,5 | 77,8 | 1,81 | 23,74774885 |
| 80 | 78 | 79 | 79,5 | 1,77 | 25,37584985 |
| 86 | 90 | 88 | 96,7 | 1,76 | 31,21771694 |

| A2_Body_fat | A2_VO2max_(ml/min/kg) | A2_VO2max_(l/min.) | A2_VT_l/min | A2_V | A2_Resting-HF |
|---|---|---|---|---|---|
| 28 | 29 | 2,05 | 1,205 | 9,4 | 69 |
| 22,6 | 41,3 | 3,317 | 2,271 | 13,4 | 54 |
| 27,5 | 41,3 | 3,927 | 2,548 | 13,4 | 79 |
| 26,5 | 35 | 2,858 | 1,523 | 11 | 69 |
| 25 | 34,3 | 2,326 | 1,704 | 11,8 | 48 |
| 18 | 43,3 | 3,424 | 2,755 | 14,2 | 60 |
| 20,6 | 36,3 | 3,488 | 2,405 | 11,8 | 57 |
| 29,8 | 26,7 | 2,083 | 2,301 | 9,4 | 72 |
| 23,1 | 37 | 3,342 | 1,515 | 10,2 | 50 |
| 24,8 | 24,7 | 2,137 | 2,354 | 9,4 | 61 |
| 29 | 32,7 | 2,634 | 1,439 | 12,6 | 54 |
| 27,5 | 37,3 | 2,922 | | 10,2 | 54 |
| 18,7 | 42,3 | 3,672 | 1,621 | 15 | 57 |
| 23,4 | 32 | 2,65 | 1,897 | 11,8 | 55 |
| 18,7 | 37,3 | 3,848 | 2,107 | 12,6 | 57 |
| 17,5 | 37,7 | 2,365 | 2,367 | 11,8 | 65 |
| 28 | 29 | 2,268 | 1,78 | 10,2 | 66 |
| 18,5 | 40 | 2,791 | 1,645 | 13,4 | 59 |
| 22,4 | 36,7 | 3,442 | 1,46 | 11,8 | 65 |
| 16,4 | 35 | 2,568 | 2,291 | 12,6 | 58 |
| 17,4 | 48,7 | 3,749 | 1,547 | 15,8 | 41 |
| 28,5 | 42,3 | 3,365 | 2,695 | 13,4 | 77 |
| 22,4 | 37,7 | 3,672 | 2,418 | 14,2 | 65 |

| A2_HFmax | A2_RQ | A2_bLa_1P | A2_Systolic_blood_pressure_right_auto_1._measurement | A2_Systolic_blood_pressure_right_auto_2._measurement | A2_Avg_systolic_blood_pressure_right_auto |
|---|---|---|---|---|---|
| 167 | 1,14 | 8,6 | 123 | 130 | 126,5 |
| 190 | 1,1 | 6,21 | 130 | 138 | 134 |
| 190 | 1,17 | 10,7 | 125 | 138 | 131,5 |
| 180 | 1,22 | 6,11 | 147 | 156 | 151,5 |
| 180 | 1,37 | 11,8 | 92 | 101 | 96,5 |
| 185 | 1,27 | 9,59 | 137 | 138 | 137,5 |
| 155 | 1,11 | 5,43 | 134 | 126 | 130 |
| 165 | 1,07 | 6,04 | 133 | 124 | 128,5 |
| 170 | 1,21 | 9,08 | 130 | 130 | 130 |
| 160 | 1,2 | 6,07 | 122 | 120 | 121 |
| 170 | 1,14 | 6,34 | 124 | 129 | 126,5 |
| 180 | 1,2 | 7,29 | 114 | 122 | 118 |
| 165 | 1,22 | 9,83 | 138 | 144 | 141 |
| 170 | 1,19 | 5,14 | 121 | 110 | 115,5 |
| 190 | 1,22 | 9,77 | 130 | 135 | 132,5 |
| 165 | 1,23 | 5,9 | 111 | 112 | 111,5 |
| 180 | 1,16 | 6,6 | 138 | 152 | 145 |
| 170 | 1,23 | 6,38 | 122 | 118 | 120 |
| 180 | 1,19 | 8,64 | 144 | 149 | 146,5 |
| 165 | 1,24 | 6,04 | 123 | 131 | 127 |
| 185 | 1,21 | 8,32 | 140 | 134 | 137 |
| 190 | 1,18 | 9,18 | 121 | 119 | 120 |
| 190 | 1,29 | 7,92 | 136 | 133 | 134,5 |

| A2_Diastolic_blood_pressure_right_auto_1._measurement | A2_Diastolic_blood_pressure_right_auto_2._measurement | A2_Avg_diastolic_blood_pressure_right_auto | A2_Cholesterol | A2_HDL | A2_LDL |
|---|---|---|---|---|---|
| 86 | 89 | 87,5 | 240 | 57,2 | 150 |
| 83 | 81 | 82 | 235 | 42 | 169 |
| 66 | 73 | 69,5 | 200 | 36,1 | 126 |
| 100 | 102 | 101 | NA | NA | NA |
| 65 | 68 | 66,5 | 170 | 35,5 | 132 |
| 84 | 82 | 83 | 201 | 47 | 119 |
| 82 | 84 | 83 | 228 | 44,8 | 159 |
| 81 | 92 | 86,5 | 255 | 52,2 | 174 |
| 79 | 80 | 79,5 | 217 | 55 | 156 |
| 79 | 82 | 80,5 | 233 | 100 | 112 |
| 78 | 80 | 79 | 206 | 85,9 | 112 |
| 74 | 81 | 77,5 | 152 | 29,5 | 108 |
| 95 | 90 | 92,5 | 197 | 38,1 | 147 |
| 82 | 78 | 80 | 229 | 47,6 | 129 |
| 79 | 82 | 80,5 | 192 | 45,1 | 132 |
| 81 | 74 | 77,5 | 197 | 75,4 | 106 |
| 94 | 100 | 97 | 247 | 80,6 | 44 |
| 71 | 68 | 69,5 | 234 | 48,8 | 157 |
| 86 | 90 | 88 | 273 | 43,9 | 164,6 |
| 78 | 86 | 82 | 196 | 31,5 | 131,6 |
| 90 | 88 | 89 | 167 | 79 | 73,9 |
| 89 | 80 | 84,5 | 179 | 54,4 | 110 |
| 86 | 90 | 88 | 188 | 29,5 | 127 |

| A2_Triglycerides | A2_Glucose | A2_uric_acid | A2_Ferritin | A2_GOT | A2_GPT |
|---|---|---|---|---|---|
| 233 | 96 | 4,2 | 19,2 | 9 | 13 |
| 97 | 95 | 4,6 | 75,7 | 20 | 21 |
| 216 | 97 | 8,5 | 105 | 15 | 28 |
| NA | NA | NA | NA | NA | NA |
| 54 | 93 | 5,6 | 34,3 | 19 | 20 |
| 159 | 106 | 4,6 | 263 | 23 | 47 |
| 118 | 94 | 7,1 | 293 | 19 | 26 |
| 176 | 104 | 5,9 | 58,5 | 13 | 21 |
| 54 | 97 | 7,4 | 248 | 12 | 46 |
| 76 | 93 | 4,6 | 7,1 | 9 | 15 |
| 46 | 96 | 5,9 | 8,5 | 10 | 20 |
| 46 | 92 | 4,1 | 8,9 | 32 | 43 |
| 81 | 96 | 5,7 | 92 | 20 | 37 |
| 182 | 89 | 6,2 | 183 | 25 | 53 |
| 67 | 94 | 6,1 | 120 | 11 | 21 |
| 49 | 78 | 4,4 | 5,7 | 9 | 14 |
| 103 | 85 | 4,8 | 17,8 | 12 | 16 |
| 156 | 103 | 5,8 | 175 | 11 | 17 |
| 407 | 87 | 5,1 | 151,5 | 17 | 29 |
| 157 | 105 | 7,9 | 217,1 | 21 | 25 |
| 41 | 102 | 4,5 | 96,3 | 17 | 31 |
| 75 | 99 | 4 | 15,3 | 13 | 20 |
| 158 | 102 | 6,9 | 249 | 28 | 47 |

| A2_Gamma-GT | A2_urea | A2_creatinine | A2_BSG | A2_Hb | A2_HKT |
|---|---|---|---|---|---|
| 8 | 16 | 0,65 | 17 | 12,7 | 38,1 |
| 20 | 35 | 0,94 | 3 | 13,4 | 41,5 |
| 34 | 38 | 0,84 | 5 | 13,1 | 39,9 |
| NA | NA | NA | NA | NA | NA |
| 15 | 22 | 0,74 | 21 | 12,2 | 37,9 |
| 19 | 45 | 0,95 | 1 | 15,3 | 44,2 |
| 18 | 49 | 0,99 | 1 | 14,6 | 42,9 |
| 26 | 22 | 0,67 | 8 | 13,9 | 42,3 |
| 131 | 35 | 0,83 | 2 | 15,2 | 45,7 |
| 15 | 27 | 0,71 | 10 | 12,3 | 36,8 |
| 11 | 37 | 0,71 | 10 | 11,6 | 37,6 |
| 18 | 28 | 0,55 | 10 | 12,6 | 38,1 |
| 145 | 40 | 0,88 | 5 | 13,6 | 39,5 |
| 52 | 35 | 0,96 | 4 | 14,9 | 45,4 |
| 25 | 28 | 0,86 | 4 | 14,8 | 42,5 |
| 13 | 28 | 0,55 | 9 | 12 | 37,8 |
| 29 | 23 | 0,65 | 5 | 11,7 | 36,7 |
| 27 | 31 | 0,82 | 3 | 13,7 | 39,6 |
| 37 | 29 | 0,9 | 3 | 14,1 | 45 |
| 18 | 41 | 0,76 | 3 | 14,2 | 42,2 |
| 37 | 37 | 0,9 | 1 | 15 | 45,6 |
| 9 | 16 | 0,57 | 2 | 13,6 | 41,2 |
| 26 | 44 | 1,09 | 8 | 15,6 | 47,4 |

| Age | Target_HF | E1_HFmax | E1_RQmax | E1_Lamax | E1_FC? |
|---|---|---|---|---|---|
| 48,93150685 | 171,0684932 | 180 | 1,17 | 8,21 | 1 |
| 46 | 174 | 200 | 1,13 | 10,2 | 1 |
| 31 | 189 | 197 | 1,29 | 10,8 | 1 |
| 54 | 166 | 180 | 1,32 | 6,64 | 1 |
| 53 | 167 | 181 | 1,3 | 9,58 | 1 |
| 37 | 183 | 175 | 1,24 | 8,2 | 1 |
| 59 | 161 | 148 | 1,07 | 3,36 | 0 |
| 57 | 163 | 170 | 1,09 | 5,55 | 0 |
| 48 | 172 | 175 | 1,26 | 8,49 | 1 |
| 58 | 162 | 166 | 1,19 | 5,29 | 1 |
| 53 | 167 | 178 | 1,19 | 6,17 | 1 |
| 40 | 180 | 190 | 1,11 | 6,55 | 1 |
| 47 | 173 | 165 | 1,16 | 10,7 | 1 |
| 62 | 158 | 180 | 1,25 | 9,29 | 1 |
| 41 | 179 | 200 | 1,26 | 6,94 | 1 |
| 43 | 177 | 190 | 1,1 | 7,96 | 1 |
| 44 | 176 | 180 | 1,29 | 6,71 | 1 |
| NA | NA | 190 | 1,24 | 5,61 | 1 |
| 46,25479452 | 173,7452055 | 185 | 1,26 | 9,05 | 1 |
| 63,03287671 | 156,9671233 | 175 | 1,32 | 6,96 | 1 |
| 52,6 | 167,4 | 184 | 1,17 | 9,18 | 1 |
| NA | NA | 190 | 1,25 | 9,02 | 1 |
| 43,9260274 | 176,0739726 | 190 | 1,29 | 11 | 1 |

| A1_HFmax | A1_RQmax | A1_Lamax | A1_FC? | E1_HFmax | E2_RQmax |
|---|---|---|---|---|---|
| 175 | 1,12 | 9,43 | 1 | 172 | 1,13 |
| 190 | 1,16 | 8,06 | 1 | 200 | 1,16 |
| 196 | 1,21 | 10,9 | 1 | 190 | 1,17 |
| 185 | 1,21 | 8,26 | 1 | 191 | 1,32 |
| 185 | 1,31 | 7,93 | 1 | 185 | 1,36 |
| 175 | 1,21 | 8,2 | 1 | 172 | 1,26 |
| 160 | 1,13 | 5,21 | 0 | 158 | 1,04 |
| 170 | 1,06 | 5,35 | 0 | 176 | 1,17 |
| 170 | 1,2 | 7,21 | 0 | 180 | 1,27 |
| 165 | 1,1 | 7,04 | 1 | 185 | 1,25 |
| 160 | 1,1 | 6,88 | 1 | 180 | 1,21 |
| 185 | 1,13 | 6,62 | 1 | 195 | 1,21 |
| 155 | 1,17 | 7,16 | 0 | 160 | 1,22 |
| 161 | 1,15 | 5,89 | 1 | 180 | 1,32 |
| 184 | 1,24 | 8,59 | 1 | 185 | 1,2 |
| 191 | 1,2 | 6,9 | 1 | 188 | 1,2 |
| 180 | 1,29 | 7,39 | 1 | 178 | 1,31 |
| 172 | 1,1 | 5,94 | 1 | 185 | 1,21 |
| 180 | 1,24 | 10,4 | 1 | 183 | 1,23 |
| 170 | 1,31 | 10,9 | 1 | 180 | 1,28 |
| 172 | 1,29 | 9,32 | 1 | 175 | 1,14 |
| 195 | 1,15 | 8,01 | 1 | 190 | 1,19 |
| 170 | 1,19 | 6,54 | 0 | 190 | 1,37 |

| E2_Lamax | E2_FC? | A2_HFmax | A2_RQmax | A2_Lamax | A2_FC? |
|---|---|---|---|---|---|
| 8,2 | 1 | 167 | 1,14 | 8,6 | 1 |
| 8,23 | 1 | 190 | 1,1 | 6,21 | 1 |
| 10,4 | 1 | 190 | 1,17 | 10,7 | 1 |
| 7,99 | 1 | 180 | 1,22 | 6,11 | 1 |
| 7,71 | 1 | 180 | 1,37 | 11,8 | 1 |
| 9,61 | 1 | 185 | 1,27 | 9,59 | 1 |
| 5,71 | 0 | 155 | 1,11 | 5,43 | 0 |
| 6,75 | 1 | 165 | 1,07 | 6,04 | 1 |
| 8,09 | 1 | 170 | 1,21 | 9,08 | 1 |
| 5,78 | 1 | 160 | 1,2 | 6,07 | 0 |
| 6,54 | 1 | 170 | 1,14 | 6,34 | 1 |
| 4,89 | 1 | 180 | 1,2 | 7,29 | 0 |
| 7,3 | 0 | 165 | 1,22 | 9,83 | 1 |
| 6,48 | 1 | 170 | 1,19 | 5,14 | 1 |
| 8,17 | 1 | 190 | 1,22 | 9,77 | 1 |
| 6,09 | 1 | 165 | 1,23 | 5,9 | 0 |
| 7,22 | 1 | 180 | 1,16 | 6,6 | 1 |
| 9,39 | 1 | 170 | 1,23 | 6,38 | 1 |
| 8,87 | 1 | 180 | 1,19 | 8,64 | 1 |
| 8,11 | 1 | 165 | 1,24 | 6,04 | 1 |
| 7,5 | 1 | 185 | 1,21 | 8,32 | 1 |
| 11,1 | 1 | 190 | 1,18 | 9,18 | 1 |
| 8,65 | 1 | 190 | 1,29 | 7,92 | 1 |

| FC_Sum |
| ---: |
| 1 |
| 1 |
| 1 |
| 1 |
| 1 |
| 1 |
| 0 |
| 0 |
| 0 |
| 0 |
| 1 |
| 0 |
| 0 |
| 1 |
| 1 |
| 0 |
| 1 |
| 1 |
| 1 |
| 1 |
| 1 |
| 1 |
| 0 |

**Table S2:** MiRNA coefficient loadings obtained with Principal Components Analysis. Higher values indicate features that explain more variance in the dimensions spanned by the Principal Components.

| miRNA | loading |
|---|---|
| hsa-miR-4665-3p | 0.126844894730938 |
| hsa-miR-940 | 0.126064037758754 |
| hsa-miR-6508-5p | 0.125611895028643 |
| hsa-miR-4649-3p | 0.124611806285172 |
| hsa-miR-6800-3p | 0.124460340421357 |
| hsa-miR-1234-3p | 0.124320811938932 |
| hsa-miR-1228-3p | 0.123791562326313 |
| hsa-miR-425-3p | 0.123544424786496 |
| hsa-miR-6889-3p | 0.123398573296676 |
| hsa-miR-6737-3p | 0.123194604578724 |
| hsa-miR-1238-3p | 0.122739133392522 |
| hsa-miR-6069 | 0.122292577917627 |
| hsa-miR-6797-3p | 0.122098652062043 |
| hsa-miR-6819-3p | 0.122054123061826 |
| hsa-miR-6813-3p | 0.121528174182025 |
| hsa-miR-6865-3p | 0.121490895026551 |
| hsa-miR-4313 | 0.121204228971606 |
| hsa-miR-1825 | 0.120596353456882 |
| hsa-miR-3162-3p | 0.120136569536415 |
| hsa-miR-4725-5p | 0.120077376077143 |
| hsa-miR-191-3p | 0.119747426978692 |
| hsa-miR-1304-3p | 0.118849711949702 |
| hsa-miR-6515-3p | 0.118392657849022 |
| hsa-miR-4787-3p | 0.114161115996535 |
| hsa-miR-1281 | 0.113985307388505 |
| hsa-miR-532-5p | 0.113130462181558 |
| hsa-miR-340-5p | 0.110701457795784 |
| hsa-miR-4433a-5p | 0.110638390740752 |
| hsa-miR-766-3p | 0.109928163884405 |
| hsa-miR-331-3p | 0.108908488711005 |
| hsa-miR-937-5p | 0.107959673181703 |
| hsa-miR-6073 | 0.10790714339371 |
| hsa-miR-150-5p | 0.107017021131117 |
| hsa-miR-93-5p | 0.106868898057692 |
| hsa-miR-342-3p | 0.106536374268913 |
| hsa-miR-532-3p | 0.106169532385253 |
| hsa-miR-378d | 0.105461057242122 |
| hsa-miR-500a-5p | 0.105367530532316 |

| | |
|---|---|
| hsa-miR-550a-3p | 0.105313116961722 |
| hsa-miR-484 | 0.1051592420621 |
| hsa-miR-361-3p | 0.104887319711668 |
| hsa-miR-30c-5p | 0.104640522533355 |
| hsa-miR-18b-5p | 0.104344224963402 |
| hsa-miR-425-5p | 0.103661256908277 |
| hsa-miR-378a-5p | 0.103543733479982 |
| hsa-miR-361-5p | 0.103243568170588 |
| hsa-let-7g-5p | 0.102716551696932 |
| hsa-miR-3195 | 0.102297327859691 |
| hsa-miR-5581-5p | 0.101927848809637 |
| hsa-miR-93-3p | 0.101740829686398 |
| hsa-miR-30d-5p | 0.10170262621957 |
| hsa-miR-26b-5p | 0.101666069611997 |
| hsa-miR-30a-5p | 0.101642615735987 |
| hsa-let-7i-5p | 0.101507048820713 |
| hsa-miR-18a-5p | 0.101499708187518 |
| hsa-miR-20a-5p | 0.101408344457159 |
| hsa-miR-17-5p | 0.101095524740345 |
| hsa-miR-20b-5p | 0.100992876749565 |
| hsa-miR-627-5p | 0.100951329719114 |
| hsa-let-7a-5p | 0.100875746752853 |
| hsa-miR-6126 | 0.100811633487655 |
| hsa-miR-144-5p | 0.100665345855181 |
| hsa-miR-17-3p | 0.100406394560773 |
| hsa-miR-15a-5p | 0.100377392827682 |
| hsa-miR-942-5p | 0.100263024260895 |
| hsa-let-7f-5p | 0.0998790815086281 |
| hsa-miR-30b-5p | 0.0997472623265359 |
| hsa-miR-128-3p | 0.0997277004844694 |
| hsa-miR-140-3p | 0.0993974869621841 |
| hsa-miR-221-3p | 0.0992238869459264 |
| hsa-miR-144-3p | 0.0990536843726823 |
| hsa-miR-6723-5p | 0.098136651486801 |
| hsa-miR-326 | 0.0981275384336112 |
| hsa-miR-6734-5p | 0.0980078710639181 |
| hsa-miR-211-3p | 0.0979576026446059 |
| hsa-miR-98-5p | 0.0976529125414225 |
| hsa-miR-4323 | 0.0972519170137695 |
| hsa-miR-3653-3p | 0.0966553056360856 |
| hsa-miR-4716-3p | 0.0966552074832214 |
| hsa-miR-340-3p | 0.0963414935269372 |
| hsa-miR-151a-3p | 0.0959178498203894 |

| | |
|---|---|
| hsa-miR-223-3p | 0.0957127427861624 |
| hsa-miR-126-3p | 0.0956141350525446 |
| hsa-miR-4306 | 0.0948196545497219 |
| hsa-miR-215-5p | 0.0948002188456182 |
| hsa-miR-500b-5p | 0.0945645308960369 |
| hsa-miR-3907 | 0.0945554101424075 |
| hsa-miR-199a-3p | 0.0945109496982375 |
| hsa-miR-16-5p | 0.0941896908331753 |
| hsa-miR-584-5p | 0.093951644221775 |
| hsa-miR-192-5p | 0.0938695486766736 |
| hsa-miR-501-3p | 0.0938396009284573 |
| hsa-miR-101-3p | 0.0935971376356248 |
| hsa-let-7d-5p | 0.0933503180262464 |
| hsa-miR-3651 | 0.0930516271962094 |
| hsa-miR-23a-3p | 0.0929451771242413 |
| hsa-miR-195-5p | 0.0928848093463347 |
| hsa-miR-199a-5p | 0.0925020329052374 |
| hsa-miR-181b-5p | 0.0921807403955276 |
| hsa-miR-186-5p | 0.0920744128760887 |
| hsa-miR-33b-3p | 0.0920672331266354 |
| hsa-miR-664a-3p | 0.092046310418814 |
| hsa-miR-29c-5p | 0.0914690994183783 |
| hsa-miR-23b-3p | 0.0911610545692254 |
| hsa-miR-328-3p | 0.0902323933404988 |
| hsa-miR-4291 | 0.0898416098831594 |
| hsa-miR-339-5p | 0.0897991325220764 |
| hsa-miR-151a-5p | 0.0896027372593557 |
| hsa-miR-454-3p | 0.089296105304483 |
| hsa-miR-8069 | 0.0891015319209009 |
| hsa-miR-939-5p | 0.0888168315698037 |
| hsa-miR-500a-3p | 0.0888041777374212 |
| hsa-miR-191-5p | 0.088732361241562 |
| hsa-miR-185-5p | 0.0881352299059626 |
| hsa-miR-197-3p | 0.0881305297314796 |
| hsa-miR-25-3p | 0.0880578987729441 |
| hsa-let-7e-5p | 0.087471626824155 |
| hsa-miR-454-5p | 0.0874027584625449 |
| hsa-miR-6724-5p | 0.0872511808332724 |
| hsa-miR-126-5p | 0.0867502997571766 |
| hsa-miR-92a-3p | 0.0859544403923714 |
| hsa-miR-21-5p | 0.0859235668509572 |
| hsa-miR-29b-3p | 0.0855004974405308 |
| hsa-miR-7114-5p | 0.0854846464741553 |

| | |
|---|---|
| hsa-miR-664b-3p | 0.0853220344375805 |
| hsa-miR-132-3p | 0.0848008333453453 |
| hsa-miR-140-5p | 0.0840632256013967 |
| hsa-miR-19b-3p | 0.084054228068963 |
| hsa-miR-6131 | 0.0836353778694813 |
| hsa-miR-196b-5p | 0.0831305588560442 |
| hsa-miR-301a-3p | 0.0830837646254873 |
| hsa-miR-151b | 0.0829276781600588 |
| hsa-miR-3135b | 0.0826969293171609 |
| hsa-miR-4732-5p | 0.0826440832272178 |
| hsa-miR-181a-5p | 0.0820045786164426 |
| hsa-miR-7152-3p | 0.0816175195335001 |
| hsa-miR-3198 | 0.0814948510483952 |
| hsa-miR-1305 | 0.0814136170472611 |
| hsa-miR-362-5p | 0.081348923606846 |
| hsa-let-7c-5p | 0.0813063806525242 |
| hsa-miR-4732-3p | 0.0812583730717475 |
| hsa-miR-4687-3p | 0.0812498397623623 |
| hsa-miR-486-5p | 0.0812220504905359 |
| hsa-miR-6826-5p | 0.0808045326721316 |
| hsa-miR-4515 | 0.0803019316294528 |
| hsa-miR-629-5p | 0.0801575991929748 |
| hsa-miR-378a-3p | 0.0797183626295239 |
| hsa-miR-4443 | 0.0792557121290484 |
| hsa-miR-505-3p | 0.0792079479492019 |
| hsa-miR-145-5p | 0.0792024419383706 |
| hsa-miR-362-3p | 0.0788494792717199 |
| hsa-miR-130a-3p | 0.0788471032717532 |
| hsa-miR-324-3p | 0.0786208862025096 |
| hsa-miR-106b-5p | 0.0783262022097773 |
| hsa-miR-365a-3p | 0.0780355816165246 |
| hsa-miR-7-1-3p | 0.0777802518818946 |
| hsa-miR-30e-5p | 0.0775960464229121 |
| hsa-miR-103a-3p | 0.0774951875803902 |
| hsa-miR-4787-5p | 0.0772216325781902 |
| hsa-miR-590-5p | 0.077053804097401 |
| hsa-miR-363-3p | 0.0770435549209454 |
| hsa-miR-96-5p | 0.0767933524546594 |
| hsa-miR-4685-5p | 0.0767724310370759 |
| hsa-miR-374b-5p | 0.0766519825277468 |
| hsa-miR-6717-5p | 0.0764455814667989 |
| hsa-miR-194-5p | 0.0763634050067215 |
| hsa-miR-130b-3p | 0.0759632240733676 |

| | |
|---|---|
| hsa-miR-148a-3p | 0.0758995592492024 |
| hsa-miR-324-5p | 0.0748805385757022 |
| hsa-miR-107 | 0.0746560735330229 |
| hsa-miR-1914-3p | 0.0746363185378495 |
| hsa-miR-15b-5p | 0.0741531420054285 |
| hsa-miR-7977 | 0.073957368038714 |
| hsa-miR-4466 | 0.0737907469546127 |
| hsa-miR-4318 | 0.0736110400863083 |
| hsa-miR-99b-5p | 0.0727522597033512 |
| hsa-let-7b-5p | 0.072719420162632 |
| hsa-miR-16-2-3p | 0.0724829340594679 |
| hsa-miR-502-3p | 0.0720831744968939 |
| hsa-miR-7-5p | 0.0720730354087346 |
| hsa-miR-210-3p | 0.0719788563806684 |
| hsa-miR-27b-3p | 0.0715241488091907 |
| hsa-miR-5100 | 0.0710660339700759 |
| hsa-miR-6780b-5p | 0.0709109084287797 |
| hsa-miR-378i | 0.0704518610621142 |
| hsa-miR-146b-5p | 0.07032301213941 |
| hsa-miR-4299 | 0.0697445454681993 |
| hsa-miR-29c-3p | 0.069638994979832 |
| hsa-miR-3679-5p | 0.0688292725078368 |
| hsa-miR-4763-3p | 0.0688204099165594 |
| hsa-miR-4713-3p | 0.068624782613144 |
| hsa-miR-296-5p | 0.0681779068605566 |
| hsa-miR-3665 | 0.0680830076881637 |
| hsa-miR-7641 | 0.06763453197541 |
| hsa-miR-6124 | 0.067208043810188 |
| hsa-miR-374a-5p | 0.0668712147478066 |
| hsa-miR-24-3p | 0.066861633772655 |
| hsa-miR-423-3p | 0.0666940246192085 |
| hsa-miR-125a-5p | 0.0663732259760776 |
| hsa-miR-4442 | 0.065399680850831 |
| hsa-miR-338-3p | 0.0650451153480166 |
| hsa-miR-4707-3p | 0.0649536583645001 |
| hsa-miR-7847-3p | 0.0646028361659879 |
| hsa-miR-183-5p | 0.064304181107325 |
| hsa-miR-624-5p | 0.0640931020410006 |
| hsa-miR-502-5p | 0.0640064453567393 |
| hsa-miR-1225-5p | 0.0636328628855675 |
| hsa-miR-99a-5p | 0.0634696240381389 |
| hsa-miR-4653-3p | 0.0630332054275075 |

| | |
|---|---|
| hsa-miR-15b-3p | 0.0623912743485709 |
| hsa-miR-1285-3p | 0.0605251385045947 |
| hsa-miR-30e-3p | 0.060514155617205 |
| hsa-miR-4459 | 0.0604659658573045 |
| hsa-miR-7107-5p | 0.0601736680238011 |
| hsa-miR-6803-3p | 0.0601214347656965 |
| hsa-miR-320a | 0.0598287941877973 |
| hsa-miR-4284 | 0.0595820614412626 |
| hsa-miR-6789-5p | 0.0592261358410231 |
| hsa-miR-1260b | 0.0590878832332045 |
| hsa-miR-574-3p | 0.0590542588757455 |
| hsa-miR-423-5p | 0.0586314840361174 |
| hsa-miR-1587 | 0.058459673705195 |
| hsa-miR-4281 | 0.0584132607312652 |
| hsa-miR-6089 | 0.0577615595359097 |
| hsa-miR-5739 | 0.057670494613895 |
| hsa-miR-1915-3p | 0.0573715378542505 |
| hsa-miR-3200-5p | 0.056022865625855 |
| hsa-miR-1207-5p | 0.0559987065398361 |
| hsa-miR-5006-5p | 0.0558337332238611 |
| hsa-miR-148b-3p | 0.0553546801299542 |
| hsa-miR-3960 | 0.0545246939772369 |
| hsa-miR-6125 | 0.0544376686582061 |
| hsa-miR-3667-5p | 0.0543480165787481 |
| hsa-miR-4286 | 0.0541989667458849 |
| hsa-miR-142-3p | 0.0541036150746894 |
| hsa-miR-222-3p | 0.0539729932724626 |
| hsa-miR-409-3p | 0.0538422302906469 |
| hsa-miR-125b-5p | 0.0535679140135874 |
| hsa-miR-6087 | 0.0534011173240026 |
| hsa-miR-342-5p | 0.052898647013272 |
| hsa-miR-7975 | 0.0527543972444341 |
| hsa-miR-1260a | 0.052550530163743 |
| hsa-miR-1268a | 0.0525004115586056 |
| hsa-miR-1246 | 0.0523936878342114 |
| hsa-miR-6085 | 0.0523896317965945 |
| hsa-miR-28-5p | 0.051770207489554 |
| hsa-miR-155-5p | 0.0513843336064061 |
| hsa-miR-4485-5p | 0.0511560653292068 |
| hsa-miR-6740-5p | 0.0510272835661965 |
| hsa-miR-4672 | 0.050799681234364 |
| hsa-miR-27a-3p | 0.0506745744512384 |
| hsa-miR-320b | 0.0504161909086582 |

| | |
|---|---|
| hsa-miR-320d | 0.0503319299749112 |
| hsa-miR-550b-2-5p | 0.0495545697569405 |
| hsa-miR-638 | 0.0486885921132907 |
| hsa-miR-4505 | 0.0478700240494632 |
| hsa-miR-625-5p | 0.047342437026334 |
| hsa-miR-6869-5p | 0.0456855275941764 |
| hsa-miR-146a-5p | 0.0450861611090543 |
| hsa-miR-744-5p | 0.0450198119095351 |
| hsa-miR-22-3p | 0.0449294318449506 |
| hsa-miR-6785-5p | 0.0448692173946502 |
| hsa-miR-574-5p | 0.044773953365457 |
| hsa-miR-660-5p | 0.0446557361367907 |
| hsa-miR-550a-3-5p | 0.0445759661237525 |
| hsa-miR-6749-5p | 0.0436427494172527 |
| hsa-miR-29a-3p | 0.043449755995708 |
| hsa-miR-4507 | 0.0432624819025495 |
| hsa-miR-320c | 0.0432073185477874 |
| hsa-miR-320e | 0.0425644347709177 |
| hsa-miR-3656 | 0.042186303039097 |
| hsa-miR-494-3p | 0.0420600563134016 |
| hsa-miR-1268b | 0.0412308319972636 |
| hsa-miR-3180-3p | 0.0399750274446369 |
| hsa-miR-6803-5p | 0.0397331383514073 |
| hsa-miR-424-5p | 0.0397303797093019 |
| hsa-miR-7704 | 0.0392112428109585 |
| hsa-miR-652-3p | 0.0389657049879776 |
| hsa-miR-8485 | 0.0381808459751307 |
| hsa-miR-100-5p | 0.0380717233480257 |
| hsa-miR-374c-5p | 0.0377156941963813 |
| hsa-miR-6875-5p | 0.0371604944267423 |
| hsa-miR-501-5p | 0.0368627629750615 |
| hsa-miR-1275 | 0.0363621512391194 |
| hsa-miR-2861 | 0.0349965794812654 |
| hsa-miR-142-5p | 0.0345845669089284 |
| hsa-miR-6088 | 0.0345109324853973 |
| hsa-miR-6800-5p | 0.0330429601493694 |
| hsa-miR-6090 | 0.0304989172291967 |
| hsa-miR-4530 | 0.0289964360886065 |
| hsa-miR-3162-5p | 0.0288627910042925 |
| hsa-miR-6127 | 0.0285554211001389 |
| hsa-miR-26a-5p | 0.0272963668359221 |

| | |
|---|---|
| hsa-miR-4788 | 0.0255443486331426 |
| hsa-miR-1202 | 0.024340581747052 |
| hsa-miR-6879-5p | 0.0225585473849342 |
| hsa-miR-642a-3p | 0.0220326972085646 |
| hsa-miR-19a-3p | 0.0218865828142491 |
| hsa-miR-4465 | 0.0215086377926363 |
| hsa-miR-1273g-3p | 0.0206106426937353 |
| hsa-miR-4497 | 0.0189308298475895 |
| hsa-miR-6165 | 0.01883958797665 |
| hsa-miR-197-5p | 0.0168384204534287 |
| hsa-miR-4516 | 0.0156095970250886 |
| hsa-miR-505-5p | 0.00963409731069468 |
| hsa-miR-6821-5p | 0.00582042707166677 |
| hsa-miR-182-5p | 0.00478000666770538 |

**Table S3:** MiRNA cluster assignments defined by cutting the hierarchical clustering dendrogram of expression values into six distinct sub-trees.

| miRNA | cluster |
|---|---|
| hsa-miR-6879-5p | 1 |
| hsa-miR-1914-3p | 1 |
| hsa-miR-6749-5p | 2 |
| hsa-miR-33b-3p | 3 |
| hsa-miR-151b | 4 |
| hsa-miR-4497 | 1 |
| hsa-miR-652-3p | 1 |
| hsa-miR-638 | 1 |
| hsa-let-7g-5p | 5 |
| hsa-miR-18b-5p | 5 |
| hsa-miR-454-3p | 5 |
| hsa-miR-939-5p | 2 |
| hsa-miR-4716-3p | 1 |
| hsa-miR-215-5p | 4 |
| hsa-miR-6785-5p | 3 |
| hsa-miR-29c-3p | 5 |
| hsa-miR-140-5p | 5 |
| hsa-let-7d-5p | 5 |
| hsa-miR-454-5p | 4 |
| hsa-miR-125a-5p | 4 |
| hsa-miR-338-3p | 5 |
| hsa-miR-23a-3p | 4 |
| hsa-miR-7-1-3p | 4 |
| hsa-miR-3200-5p | 6 |
| hsa-miR-374a-5p | 5 |
| hsa-miR-362-5p | 6 |

| | |
|---|---|
| hsa-miR-30a-5p | 2 |
| hsa-miR-151a-5p | 4 |
| hsa-miR-6737-3p | 3 |
| hsa-miR-18a-5p | 5 |
| hsa-miR-6088 | 3 |
| hsa-miR-6089 | 3 |
| hsa-miR-6087 | 3 |
| hsa-miR-6085 | 2 |
| hsa-miR-16-5p | 5 |
| hsa-miR-4530 | 3 |
| hsa-miR-185-5p | 2 |
| hsa-miR-6813-3p | 3 |
| hsa-miR-342-5p | 1 |
| hsa-miR-374c-5p | 5 |
| hsa-miR-4665-3p | 3 |
| hsa-miR-186-5p | 2 |
| hsa-miR-378a-5p | 4 |
| hsa-miR-125b-5p | 4 |
| hsa-miR-3665 | 1 |
| hsa-miR-766-3p | 4 |
| hsa-miR-664b-3p | 4 |
| hsa-miR-664a-3p | 4 |
| hsa-miR-6090 | 3 |
| hsa-miR-6508-5p | 3 |
| hsa-miR-100-5p | 4 |
| hsa-let-7e-5p | 5 |
| hsa-miR-320e | 2 |
| hsa-miR-1225-5p | 3 |
| hsa-miR-19b-3p | 4 |
| hsa-miR-27b-3p | 5 |
| hsa-miR-197-5p | 3 |
| hsa-miR-1228-3p | 3 |
| hsa-miR-140-3p | 4 |
| hsa-miR-211-3p | 2 |
| hsa-miR-196b-5p | 5 |
| hsa-miR-148a-3p | 5 |
| hsa-miR-1260b | 2 |
| hsa-miR-93-5p | 5 |
| hsa-miR-574-5p | 3 |
| hsa-miR-148b-3p | 5 |
| hsa-miR-6826-5p | 2 |
| hsa-miR-132-3p | 6 |
| hsa-miR-28-5p | 4 |
| hsa-miR-151a-3p | 2 |
| hsa-miR-424-5p | 5 |
| hsa-miR-7641 | 1 |
| hsa-miR-6069 | 3 |

| | |
|---|---|
| hsa-miR-4516 | 3 |
| hsa-miR-4515 | 3 |
| hsa-miR-128-3p | 4 |
| hsa-miR-1260a | 2 |
| hsa-miR-942-5p | 4 |
| hsa-miR-6740-5p | 1 |
| hsa-miR-342-3p | 4 |
| hsa-miR-1238-3p | 3 |
| hsa-miR-532-3p | 4 |
| hsa-miR-8069 | 3 |
| hsa-miR-6889-3p | 3 |
| hsa-miR-4281 | 3 |
| hsa-miR-4286 | 2 |
| hsa-miR-4284 | 3 |
| hsa-miR-30e-3p | 4 |
| hsa-miR-146a-5p | 5 |
| hsa-miR-25-3p | 2 |
| hsa-miR-7704 | 1 |
| hsa-miR-6865-3p | 3 |
| hsa-miR-223-3p | 4 |
| hsa-miR-6073 | 5 |
| hsa-miR-4505 | 1 |
| hsa-miR-4507 | 1 |
| hsa-miR-5581-5p | 1 |
| hsa-miR-324-5p | 6 |
| hsa-miR-1915-3p | 1 |
| hsa-miR-502-3p | 6 |
| hsa-miR-584-5p | 6 |
| hsa-miR-4299 | 3 |
| hsa-miR-3960 | 3 |
| hsa-miR-4291 | 5 |
| hsa-miR-328-3p | 4 |
| hsa-miR-93-3p | 4 |
| hsa-miR-363-3p | 5 |
| hsa-miR-30b-5p | 4 |
| hsa-miR-21-5p | 5 |
| hsa-miR-199a-5p | 4 |
| hsa-miR-144-5p | 5 |
| hsa-miR-145-5p | 4 |
| hsa-miR-425-5p | 4 |
| hsa-miR-409-3p | 4 |
| hsa-miR-98-5p | 5 |
| hsa-miR-3195 | 2 |
| hsa-miR-3198 | 1 |
| hsa-miR-6131 | 1 |
| hsa-miR-1587 | 1 |
| hsa-miR-23b-3p | 4 |

| | |
|---|---|
| hsa-miR-5739 | 2 |
| hsa-miR-500a-5p | 6 |
| hsa-miR-425-3p | 3 |
| hsa-miR-7977 | 2 |
| hsa-miR-146b-5p | 5 |
| hsa-miR-2861 | 1 |
| hsa-let-7b-5p | 5 |
| hsa-miR-6819-3p | 3 |
| hsa-miR-192-5p | 4 |
| hsa-miR-624-5p | 5 |
| hsa-miR-126-5p | 5 |
| hsa-miR-191-5p | 4 |
| hsa-miR-590-5p | 5 |
| hsa-miR-1825 | 3 |
| hsa-miR-24-3p | 5 |
| hsa-miR-4732-3p | 4 |
| hsa-miR-532-5p | 5 |
| hsa-miR-629-5p | 1 |
| hsa-miR-103a-3p | 5 |
| hsa-miR-6789-5p | 1 |
| hsa-miR-361-3p | 4 |
| hsa-miR-6800-3p | 3 |
| hsa-miR-15b-5p | 5 |
| hsa-miR-744-5p | 1 |
| hsa-miR-6803-3p | 4 |
| hsa-miR-324-3p | 2 |
| hsa-miR-326 | 4 |
| hsa-miR-6125 | 3 |
| hsa-miR-6124 | 3 |
| hsa-miR-6127 | 1 |
| hsa-miR-6126 | 2 |
| hsa-miR-6724-5p | 1 |
| hsa-miR-502-5p | 6 |
| hsa-miR-423-3p | 4 |
| hsa-miR-4685-5p | 1 |
| hsa-miR-3162-5p | 3 |
| hsa-let-7a-5p | 5 |
| hsa-miR-627-5p | 6 |
| hsa-miR-6875-5p | 1 |
| hsa-miR-4653-3p | 1 |
| hsa-miR-6797-3p | 3 |
| hsa-miR-181b-5p | 5 |
| hsa-miR-107 | 5 |
| hsa-miR-3907 | 2 |
| hsa-miR-4787-3p | 3 |
| hsa-let-7i-5p | 5 |
| hsa-miR-1234-3p | 3 |

| | |
|---|---|
| hsa-miR-106b-5p | 5 |
| hsa-miR-5006-5p | 3 |
| hsa-miR-221-3p | 5 |
| hsa-miR-320d | 2 |
| hsa-miR-320c | 2 |
| hsa-miR-320b | 2 |
| hsa-miR-320a | 2 |
| hsa-miR-301a-3p | 5 |
| hsa-miR-4687-3p | 1 |
| hsa-miR-5100 | 2 |
| hsa-miR-7107-5p | 3 |
| hsa-miR-501-5p | 1 |
| hsa-miR-4732-5p | 2 |
| hsa-miR-181a-5p | 5 |
| hsa-miR-1202 | 3 |
| hsa-miR-4313 | 3 |
| hsa-miR-17-3p | 5 |
| hsa-miR-4318 | 5 |
| hsa-miR-183-5p | 5 |
| hsa-let-7c-5p | 5 |
| hsa-miR-126-3p | 5 |
| hsa-miR-92a-3p | 2 |
| hsa-miR-182-5p | 4 |
| hsa-miR-4433a-5p | 3 |
| hsa-miR-30c-5p | 4 |
| hsa-miR-15a-5p | 5 |
| hsa-miR-191-3p | 3 |
| hsa-miR-29b-3p | 5 |
| hsa-miR-4788 | 1 |
| hsa-miR-6800-5p | 1 |
| hsa-miR-15b-3p | 4 |
| hsa-miR-3180-3p | 1 |
| hsa-miR-3135b | 3 |
| hsa-miR-4649-3p | 3 |
| hsa-miR-3162-3p | 3 |
| hsa-miR-937-5p | 1 |
| hsa-miR-6723-5p | 2 |
| hsa-miR-378d | 5 |
| hsa-miR-378i | 6 |
| hsa-miR-1275 | 3 |
| hsa-miR-4306 | 2 |
| hsa-miR-6869-5p | 1 |
| hsa-miR-29c-5p | 2 |
| hsa-miR-1305 | 1 |
| hsa-miR-625-5p | 1 |
| hsa-miR-1285-3p | 6 |
| hsa-miR-7-5p | 5 |

| | |
|---|---|
| hsa-miR-3656 | 1 |
| hsa-miR-3651 | 3 |
| hsa-miR-500b-5p | 6 |
| hsa-miR-550a-3p | 4 |
| hsa-miR-6734-5p | 1 |
| hsa-miR-4442 | 1 |
| hsa-miR-4443 | 1 |
| hsa-miR-4763-3p | 1 |
| hsa-miR-130a-3p | 5 |
| hsa-miR-4713-3p | 1 |
| hsa-miR-195-5p | 5 |
| hsa-miR-30e-5p | 5 |
| hsa-miR-296-5p | 4 |
| hsa-miR-505-5p | 1 |
| hsa-miR-660-5p | 5 |
| hsa-miR-3679-5p | 3 |
| hsa-miR-6717-5p | 1 |
| hsa-miR-550a-3-5p | 5 |
| hsa-miR-222-3p | 5 |
| hsa-miR-4707-3p | 4 |
| hsa-miR-19a-3p | 4 |
| hsa-miR-142-3p | 4 |
| hsa-miR-197-3p | 4 |
| hsa-miR-339-5p | 4 |
| hsa-miR-7847-3p | 1 |
| hsa-miR-331-3p | 4 |
| hsa-miR-4459 | 3 |
| hsa-miR-20b-5p | 5 |
| hsa-miR-486-5p | 4 |
| hsa-miR-101-3p | 5 |
| hsa-miR-4323 | 4 |
| hsa-miR-27a-3p | 5 |
| hsa-miR-1268b | 3 |
| hsa-miR-16-2-3p | 2 |
| hsa-miR-550b-2-5p | 5 |
| hsa-miR-574-3p | 4 |
| hsa-miR-3653-3p | 3 |
| hsa-miR-1304-3p | 3 |
| hsa-miR-4485-5p | 3 |
| hsa-miR-4466 | 1 |
| hsa-miR-4465 | 1 |
| hsa-miR-484 | 4 |
| hsa-miR-500a-3p | 6 |
| hsa-miR-642a-3p | 3 |
| hsa-miR-340-3p | 4 |
| hsa-miR-6165 | 3 |
| hsa-miR-1246 | 3 |

| | |
|---|---|
| hsa-miR-501-3p | 6 |
| hsa-miR-374b-5p | 5 |
| hsa-miR-505-3p | 4 |
| hsa-miR-494-3p | 3 |
| hsa-miR-1268a | 3 |
| hsa-miR-142-5p | 4 |
| hsa-miR-361-5p | 4 |
| hsa-miR-20a-5p | 5 |
| hsa-miR-210-3p | 6 |
| hsa-miR-96-5p | 5 |
| hsa-miR-1273g-3p | 3 |
| hsa-miR-1207-5p | 3 |
| hsa-miR-6803-5p | 1 |
| hsa-miR-17-5p | 5 |
| hsa-miR-4672 | 1 |
| hsa-miR-423-5p | 2 |
| hsa-miR-4725-5p | 3 |
| hsa-miR-99a-5p | 4 |
| hsa-miR-155-5p | 5 |
| hsa-miR-194-5p | 4 |
| hsa-miR-7975 | 2 |
| hsa-miR-940 | 3 |
| hsa-miR-365a-3p | 4 |
| hsa-miR-6821-5p | 3 |
| hsa-miR-99b-5p | 4 |
| hsa-let-7f-5p | 5 |
| hsa-miR-7152-3p | 1 |
| hsa-miR-4787-5p | 3 |
| hsa-miR-6515-3p | 3 |
| hsa-miR-362-3p | 4 |
| hsa-miR-150-5p | 4 |
| hsa-miR-130b-3p | 6 |
| hsa-miR-22-3p | 6 |
| hsa-miR-26b-5p | 5 |
| hsa-miR-199a-3p | 5 |
| hsa-miR-30d-5p | 2 |
| hsa-miR-144-3p | 5 |
| hsa-miR-26a-5p | 4 |
| hsa-miR-29a-3p | 4 |
| hsa-miR-7114-5p | 2 |
| hsa-miR-8485 | 3 |
| hsa-miR-340-5p | 6 |
| hsa-miR-6780b-5p | 1 |
| hsa-miR-378a-3p | 6 |
| hsa-miR-3667-5p | 6 |
| hsa-miR-1281 | 3 |

**Table S4:** MiRNA variable importance assigned by performing linear regression on the VO2 max levels using expression vectors as features. A higher variable importance indicates a stronger approximated effect on VO2 max levels. In particular, correlated miRNAs can share a high coefficient value.

| miRNA | variable importance |
|---|---|
| hsa-miR-4465 | 1.14996521708515 |
| hsa-miR-409-3p | 0.920982372217814 |
| hsa-miR-3135b | 0.851367880095478 |
| hsa-miR-5006-5p | 0.809146433880345 |
| hsa-miR-365a-3p | 0.77546932979795 |
| hsa-miR-340-5p | 0.751770394668099 |
| hsa-miR-7975 | 0.743552471889329 |
| hsa-miR-362-5p | 0.692950168865217 |
| hsa-miR-1202 | 0.670374971865862 |
| hsa-miR-26a-5p | 0.612990294055711 |
| hsa-let-7d-5p | 0.605533856557137 |
| hsa-miR-502-5p | 0.599264301286434 |
| hsa-miR-5581-5p | 0.495028931866778 |
| hsa-miR-642a-3p | 0.466952939424561 |
| hsa-let-7b-5p | 0.466318683577534 |
| hsa-miR-183-5p | 0.457505975197679 |
| hsa-miR-486-5p | 0.445193776320586 |
| hsa-miR-6879-5p | 0.42191555564153 |
| hsa-miR-454-5p | 0.407482186537414 |
| hsa-miR-28-5p | 0.399790201366784 |
| hsa-miR-4459 | 0.391825566593816 |
| hsa-miR-6869-5p | 0.378641319646266 |
| hsa-miR-4323 | 0.345947600257929 |
| hsa-miR-502-3p | 0.340357054638963 |
| hsa-miR-1273g-3p | 0.314232360084127 |
| hsa-miR-338-3p | 0.304834733926964 |
| hsa-miR-6821-5p | 0.293484029411793 |
| hsa-miR-505-3p | 0.290679056939262 |
| hsa-miR-222-3p | 0.274157371164798 |
| hsa-miR-126-5p | 0.270081455484812 |
| hsa-miR-1207-5p | 0.269415458231481 |
| hsa-miR-424-5p | 0.26709146896224 |
| hsa-miR-146b-5p | 0.264972262450903 |
| hsa-miR-27a-3p | 0.26289905398942 |
| hsa-let-7g-5p | 0.257663708881599 |
| hsa-miR-144-5p | 0.257658568949595 |
| hsa-miR-221-3p | 0.253559394762969 |
| hsa-miR-182-5p | 0.248513713988872 |

| | |
|---|---|
| hsa-miR-1268a | 0.247022575753072 |
| hsa-miR-3960 | 0.24629597631404 |
| hsa-miR-6819-3p | 0.246123214799376 |
| hsa-miR-4707-3p | 0.231706293215975 |
| hsa-miR-500a-3p | 0.214960777155456 |
| hsa-miR-181a-5p | 0.20280917898196 |
| hsa-miR-4466 | 0.195475109157439 |
| hsa-miR-6165 | 0.192853839470799 |
| hsa-miR-1914-3p | 0.190343193934103 |
| hsa-miR-6803-5p | 0.179571354329639 |
| hsa-miR-8069 | 0.176611282130342 |
| hsa-miR-145-5p | 0.172686258699069 |
| hsa-miR-3653-3p | 0.171825693382487 |
| hsa-miR-652-3p | 0.168227145477399 |
| hsa-let-7a-5p | 0.166664201261705 |
| hsa-miR-625-5p | 0.161640966082536 |
| hsa-miR-100-5p | 0.160833829426904 |
| hsa-miR-505-5p | 0.150845370502293 |
| hsa-miR-107 | 0.150493948280093 |
| hsa-miR-4732-3p | 0.148424431994047 |
| hsa-miR-6073 | 0.138187911973151 |
| hsa-miR-6089 | 0.108964714026655 |
| hsa-miR-4516 | 0.106137830340153 |
| hsa-miR-155-5p | 0.101677960475269 |
| hsa-miR-4653-3p | 0.0989085501142673 |
| hsa-miR-423-5p | 0.0956324481486475 |
| hsa-miR-29c-3p | 0.0953009070058705 |
| hsa-miR-7641 | 0.0948130807555747 |
| hsa-miR-4433a-5p | 0.0933749121776901 |
| hsa-miR-4286 | 0.087054445775337 |
| hsa-miR-106b-5p | 0.0719747990542692 |
| hsa-miR-361-3p | 0.0708474527094351 |
| hsa-miR-4672 | 0.0696518507109891 |
| hsa-miR-132-3p | 0.0693322409597973 |
| hsa-miR-3907 | 0.0668006372814346 |
| hsa-miR-532-5p | 0.063799722218583 |
| hsa-miR-937-5p | 0.050845881672489 |
| hsa-miR-6723-5p | 0.0490898980928391 |
| hsa-let-7i-5p | 0.0454008992844827 |
| hsa-miR-378d | 0.0441184451473226 |
| hsa-miR-29a-3p | 0.0409778151678459 |
| hsa-miR-6131 | 0.0340326642079648 |

| | |
|---|---|
| hsa-miR-7-5p | 0.0301254079679339 |
| hsa-miR-574-5p | 0.028769524888004 |
| hsa-miR-186-5p | 0.0239300296456704 |
| hsa-miR-6088 | 0.0224342283183027 |
| hsa-miR-4284 | 0.0178374950425057 |
| hsa-miR-92a-3p | 0.00846095933573645 |
